# Species traits and landscape structure can drive scale-dependent propagation of effects in ecosystems

**DOI:** 10.1101/2023.11.15.567315

**Authors:** David García-Callejas, Sandra Lavorel, Otso Ovaskainen, Duane A. Peltzer, Jason M. Tylianakis

**Affiliations:** Centre for Integrative Ecology, School of Biological Sciences, University of Canterbury, Private Bag 4800, Christchurch 8140, New Zealand.; Manaaki Whenua-Landcare Research, PO Box 69040, Lincoln 7640, New Zealand.; Laboratoire d’Ecologie Alpine, Université Grenoble Alpes, Université Savoie Mont-Blanc, CNRS, Grenoble, France; Department of Biological and Environmental Science, University of Jyväskylä, P.O. Box 35 (Survontie 9C), FI-40014 Jyväskylä, Finland; Organismal and Evolutionary Biology Research Programme, Faculty of Biological and Environmental Sciences, University of Helsinki, P.O. Box 65, Helsinki 00014, Finland; Bioprotection Aotearoa, School of Biological Sciences, University of Canterbury, Private Bag 4800, Christchurch 8140, Aotearoa New Zealand

## Abstract

Species can directly and indirectly affect others across communities and habitats, yet the spatial scale over which such effects spread remains unclear. This uncertainty arises partly because the species traits and landscape structures allowing indirect effects to propagate may differ across scales. Here, we use a topological network metric, communicability, to explore the spatial propagation of effects in a large-scale plant-frugivore network projected across the territory of Aotearoa New Zealand. We show that generalism and species prevalence, and complementary morphological traits such as fruit and body size, are important predictors of species’ capacity to propagate effects, but their importance differed across scales. Furthermore, native bird species (but not exotics) showed a positive relationship between body size and their potential to propagate effects. Habitat composition was the most important landscape factor in our study, generating hotspots of effect propagation around forested areas, whereas landscapes containing a variety of habitats acted as a buffer against propagation. Overall, our results indicate that species displaying specific sets of traits, including ubiquity, interaction generalism, and a combination of large body size and native status, are the most likely to propagate large-scale ecological impacts in the plant-frugivore communities studied, yet landscape properties may moderate this spread.

## Introduction

Species can directly influence the population dynamics of their interaction partners through consumption or facilitation, or those of non-partner species through one or more shared partners, in what are usually named *indirect effects* (Higashi & Nakajima, 1995; Holt & Bonsall, 2017; Nakajima & Higashi, 1995; Pires *et al*., 2020; Wootton, 2002). These direct and indirect effects provide the pathway for disturbances to propagate across space by altering species’ population dynamics (Martins *et al*., 2024). However, ecologists cannot currently predict the rate and extent that these indirect effects propagate in space, making it crucial to understand the relative importance of different species and habitats in connecting communities and ecosystems (Hackett *et al*., 2019; McCann *et al*., 2005; McCoy *et al*., 2009).

Measuring the spatial propagation of ecological effects across communities is challenging because any two species in natural systems can be connected via multiple direct and indirect interaction pathways, which are exceedingly difficult to track simultaneously. As a consequence, our understanding of indirect effects in natural systems is largely derived from studies where such indirect effects are linear and non-spatial, or from small, experimentally manipulable communities. Initial work on trophic cascades revealed that biomass of a primary resource is indirectly affected by a top predator, via changes in the biomass of an intermediate consumer (e.g. Ripple *et al*. (2016)). Theory (Higashi & Nakajima, 1995; Nakajima & Higashi, 1995), dynamical simulations of interaction networks (García-Callejas *et al*., 2019; Montoya *et al*., 2009), and experimental species removals from communities (Donohue *et al*., 2017) demonstrated the importance of indirect interactions for both population (Wootton, 2002) and evolutionary (Cosmo *et al*., 2023; Guimarães Jr *et al*., 2017) processes. Additional studies combined network analysis with experimental manipulations to demonstrate indirect effects among hosts via shared parasitoids (Morris *et al*., 2004; Tack *et al*., 2011). This body of literature demonstrates that the impacts of indirect effects in communities are comparable to, and sometimes greater than, those of direct effects.

In contrast to these indirect effects within single, discrete communities of interacting species, populations and communities are open systems in which spatial fluxes of individuals and nutrients underpin their dynamics (Gounand *et al*., 2018; Gravel *et al*., 2016). The spatial signature of indirect effects is therefore fundamental for understanding community dynamics. Yet, empirical interaction networks are typically generated by aggregating data in space (and time), such that nodes represent species that lack any specific spatial location. Metacommunity and metanetwork studies capture spatial processes such as dispersal, but even these treat individual patches as discrete, albeit connected, interacting communities (Gounand *et al*., 2018; Jacobson & Peres-Neto, 2010). Nevertheless, such metanetwork studies have revealed cross-habitat indirect effects (Frost *et al*., 2016; Harvey *et al*., 2016; Knight *et al*., 2005), whereby fluxes of nutrients across habitats can influence the dynamics of (Wootton *et al*., 2023), and even destabilise (McCann *et al*., 2021; Subalusky & Post, 2019), the recipient habitat. Likewise, the movement of individuals may induce complex dynamics and feedbacks between habitats, e.g. via increased predation in one habitat (McCoy *et al*., 2009) or coupling through apparent competition (Frost *et al*., 2016; Holt & Bonsall, 2017), and the indirect effects mediated by species movement potentially differ from spatial subsidies of nutrients (Wootton *et al*., 2023). Despite these fundamental insights from spatially-coupled habitat patches, mainland populations of species are distributed continuously in space, such that their movement and interactions should indirectly connect the dynamics of populations across a range of scales. However, the spatial patterns of indirect effects have not been quantified empirically at larger scales or in more complex settings, and the role of different species or spatial configurations in shaping such spatially-explicit indirect effects remains unclear (Martins *et al*., 2024). Thus, we lack the conceptual ground on which to build theory and models, the quantitative metrics with which to capture how effects propagate across ecosystems, and the large-scale spatial data with which to test and refine hypotheses.

Species’ propensity to spatially propagate indirect effects depends on their interactions with other species, their movement capacities, behavioural traits, and abundance (Martins *et al*., 2024). First, species’ traits can predict their tendency to interact with others (Bartomeus *et al*., 2016) and to link spatial locations through dispersal (Raffard *et al*., 2022). Second, although species’ tendency to connect with others within a habitat can vary independently of their role in connecting across habitats (Hackett *et al*., 2019), traits such as trophic generalism (within a habitat) can determine a species’ propensity to spill over from one habitat to another (Frost *et al*., 2015). Landscape structure will, in turn, influence species movement within and across habitats (Boesing *et al*., 2018). For example, areas holding high species diversity and abundance may act as source areas for nearby habitats, facilitating the spatial propagation of indirect effects.

Here we assess the relative importance of species traits and landscape variables for determining the scale over which biotic indirect effects propagate through space. We intentionally refer to the propagation of *effects*, as that term encompasses any potential processes propagating between any two nodes in a network. Our network approach assumes that direct interspecific interactions modify species densities, and species that are not directly connected can affect each other through “interaction chains”, i.e. series of direct interactions between pairs of species that modify their densities (Wootton, 2002), also called “density-mediated indirect interactions” (Abrams *et al*., 1996). To facilitate understanding of such effects from small to large scales, we first introduce to ecology a metric from network theory to quantify propagation across all direct and indirect paths between species. This metric, *communicability*, estimates the propagation of effects between any two nodes in a network only from information about its structure (Estrada & Hatano, 2008), potentially overcoming the need to quantify the population dynamics of the interacting nodes (i.e. species). We begin by demonstrating the link between this topological metric and community dynamics.

Given the definition of communicability (see Methods) and previous work on the importance of highly-connected species in driving different ecological patterns (e.g. neutral interactions (Peralta *et al*., 2020), metacommunity complexity and diversity (Pillai *et al*., 2011), habitat coupling (Frost *et al*., 2015; Kortsch *et al*., 2015)), we hypothesise that the propensity to propagate effects in space will be higher in 1) species with a high generalism (e.g. high network degree), and 2) at larger spatial scales, in species with a wide distribution, which are able to reach larger parts of a given territory. Although we expect generalist species to have the most paths for influencing others, degree does not necessarily predict communicability (Estrada & Hatano, 2008), particularly in nested networks (Jonhson *et al*., 2013) - an architecture that is typical of mutualistic interactions (Mariani *et al*., 2019).

Based on these first-principles, we further hypothesise that traits (morphological, behavioural, or otherwise) correlated with interaction generalism and prevalence will impact the propensity of species to indirectly affect many others across space (Martins *et al*., 2024). For example, exotic species are often more generalist than natives (Aizen *et al*., 2008), which can also allow them to live in a greater range of habitats (Marvier *et al*., 2004). We therefore expect provenance to be a key trait determining species’ propensity to propagate effects in space. In contrast, focusing on plant-frugivore interactions, larger fruits can only be swallowed by large birds (potentially leading to lower partner generalism in large-fruited species), yet large birds may disperse seeds further (Wotton & Kelly, 2012), influencing a plant’s distribution. At larger scales, we hypothesise that habitats that tend to harbour a high diversity of native bird and fruiting plant species, such as forest and shrubland, will be the most important for propagating effects. In contrast, landscapes with many habitat boundaries (i.e. high habitat diversity) may slow the propagation of effects, as species turnover across habitats will lengthen the indirect pathways needed to traverse space.

To test the fundamental principles behind the spatial spread of indirect effects and address these knowledge gaps, we analyze how effects are propagated in a large-scale ecological system: the network of plants and bird frugivores across the territory of Aotearoa New Zealand. Although datasets of empirical networks across a range of sites exist, empirical studies typically aim for spatial independence among replicate networks, and their spacing is typically arbitrary. In contrast, understanding the spatial propagation of indirect effects requires a continuous ‘layer’ of connections at different scales. We therefore overcome data limitations by generating a data-informed country-scale spatial network of species interactions, considering both native and exotic species. Through this approach, we evaluate which traits (provenance, morphological or interaction traits) explain species’ capacity to propagate effects across scales, and which landscape properties promote or hinder the propagation of effects across the territory.

## Results

### Overview: Large-scale frugivory network and spatial indirect effects

To explore systematically the propagation of effects across space, we projected the occurrences and interactions of 102 plant species and 22 frugivore birds across the two main islands of Aotearoa New Zealand. For building the spatial country-scale frugivory network (Fig. 1), we combined a) species occurrences, modelled with a spatially-explicit Joint Species Distribution Model (Tikhonov *et al*., 2020b), b) a metaweb, i.e. the set of all known frugivory interactions between plants and birds compiled from the literature (Peralta *et al*., 2020), and c) bird dispersal kernels, estimated from their Hand-Wing Index (Sheard *et al*., 2020), to determine the limits of spatial connections among populations. We then obtained the communicability measure of every population in each cell. For any given species *a*, we refer to a local node of that species in a cell *i*, as a population of that species (e.g. *a_i_* in panel D of Fig. 1). Finally, we related a series of species-level traits and landscape factors to population-level communicability values (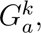 see Methods), aggregated species-level communicability across all the territory (*S_G__a_*), and landscape-level communicability (e.g. the sum of all communicability values, across all species, in a given cell, *L_G_^k^*), using generalised linear models.

**Fig. 1:**
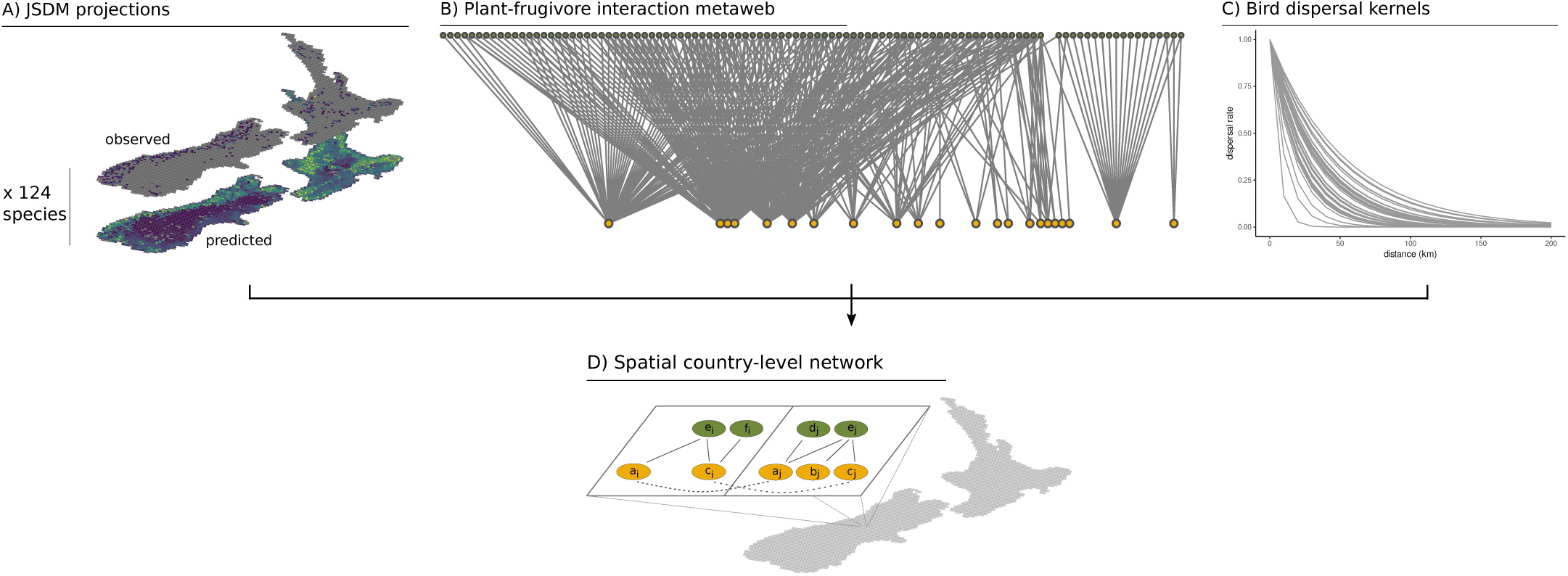
Visual summary of the process for building a large-scale spatial network of plant-frugivore interactions. In A) we projected the occurrences of 22 bird species and 102 plant species over the two main islands of Aotearoa New Zealand, using empirical data from systematic surveys and a joint species distribution model (see Methods). In B) we limited each species’ potential range of partners using the set of recorded interactions between these species, i.e. the metaweb, from Peralta *et al*. (2020). In C) we estimated dispersal kernels for bird species using their Hand-Wing Index as a proxy (Sheard *et al*., 2020). From these steps, we projected bird and plant species occurrences, added their known interactions from the metaweb, and connected populations via bird dispersal in a grid of 3030 cells of 10x10km. In D), two adjacent cells are depicted. Each cell holds interactions between the locally present species (continous links), and potential bird dispersal to other cells (dashed links), depending on the dispersal capacities of the species present. which were predicted to be present in more than two thirds of the territory (*>* 2000 cells out of 3030).

### Characteristics of the Aotearoa New Zealand plant-frugivore network

The metaweb of plant-frugivore interactions in Aotearoa New Zealand, after filtering for those species with available trait data, is composed of 102 plant species and 22 bird species, of which 92 plants and 14 birds are native. It has a connectance of 0.44, and the most generalist species are the New Zealand bellbird (*Anthornis melanura*), the kererū (*Hemiphaga novaeseelandiae*), and the silvereye (*Zosterops lateralis*) with 57 interaction partners recorded for each species, while there are 27 plant species with only one recorded bird frugivore in the metaweb. The most prevalent bird species according to our distribution analysis were a mixture of exotics (the eurasian chaffinch (*Fringilla coelebs*, Fringillidae) and the blackbird (*Turdus merula*, Turdidae)) and natives (grey warbler (*Gerygone igata*, Acanthizidae)) and silvereye (*Zosterops lateralis*, Zosteropidae)), all of which were predicted to be present in more than two thirds of the territory (> 2000 cells out of 3030).

### Population and species-level communicability

We found that the communicability metric presented here shows a good agreement with dynamic estimates of net effects between species in mutualistic systems (measured using established methods, as in e.g Higashi & Nakajima (1995); Montoya *et al*. (2009); Novak *et al*. (2016)). We therefore consider it a valid first approximation of the potential to propagate direct and indirect effects between populations, in the absence of detailed empirical information. This is explored in the Supplementary Section “Structural and dynamical propagation”, where we show that the structural and dynamical estimates of indirect effects are generally well correlated in competitive and mutualistic networks, but not in food webs, which include both positive (for consumers) and negative (for resources) effects.

In the Aotearoa New Zealand plant-frugivore network, the ranking of species according to their communicability varied depending on the spatial scale considered. Within local communities (panel A of Fig. 2), the kererū (*Hemiphaga novaeseelandiae*, a native bird from the Columbidae family) showed the highest average population-level communicability (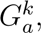 eq. 3), followed by raukawa (*Raukaua edgerleyi*), a native plant from the Araliaceae family. However, the most important species when aggregating all local populations, i.e. at the scale of the whole study area (*S_G__a_*, eq. 4, panel B of Fig. 2), were two bird species with widespread distributions in Aotearoa New Zealand: the silvereye and the blackbird. Silvereyes were self-introduced from Australia in the first half of the 19th century, so here we consider them a native species. Blackbirds, on the other hand, are a worldwide distributed exotic species.

**Fig. 2:**
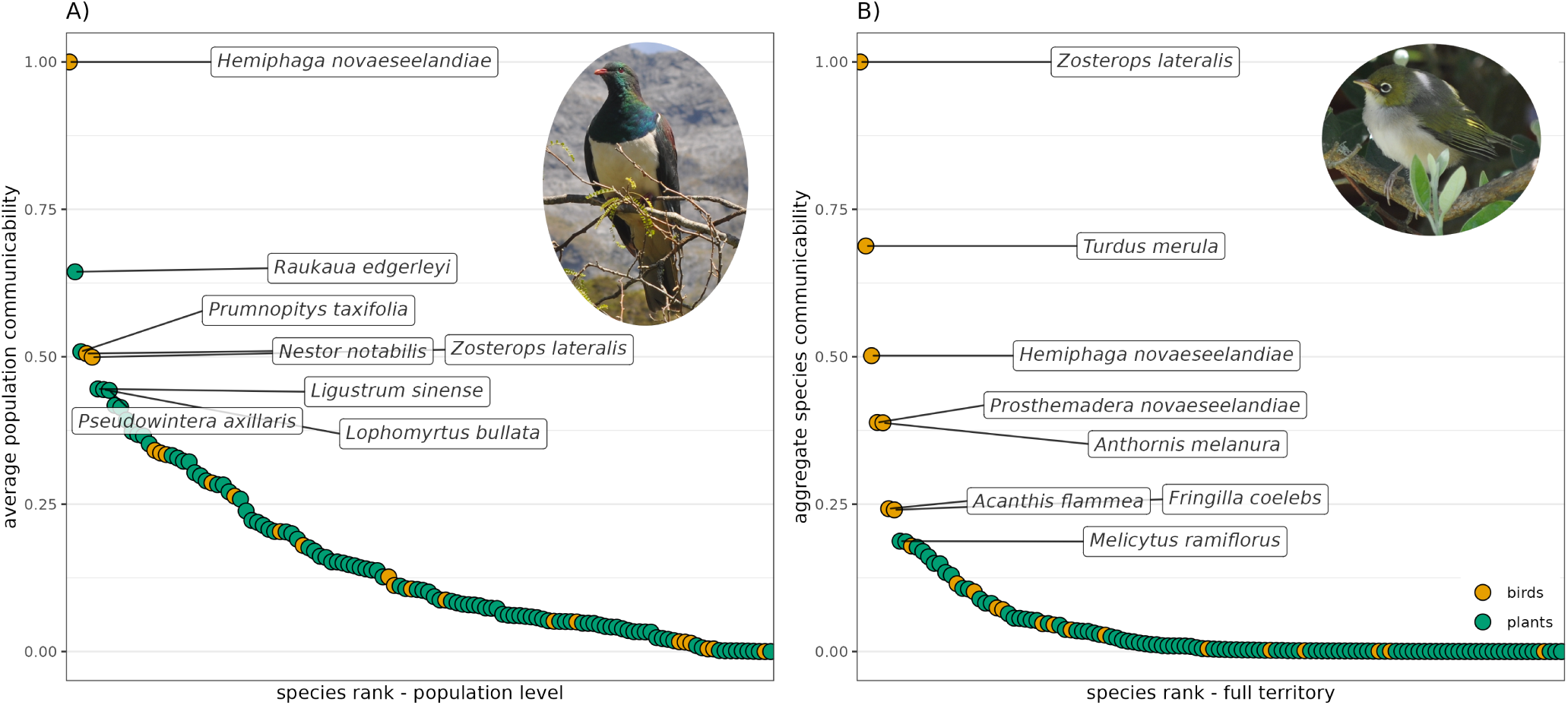
Relative communicability values at the scale of local populations (panel A) and the whole study area (panel B). Communicability values are scaled in the range 0-1 in both cases. Inset pictures show, on the left panel, the kererū (*Hemiphaga novaeseelandiae*), and on the right panel, the silvereye (*Zosterops lateralis*). Photos by Lucas P. Martins.

Shifts in relative communicability rankings across scales emerged from the interplay between generalism (degree) and prevalence. At the country level, species prevalence and degree significantly influenced species-level communicability in plants and birds, as expected, but species provenance and morphological traits were also relevant predictors. Regarding plants, native species showed a higher species-level communicability. The interaction between fruit size and provenance was also statistically significant, showing that the effect of fruit size on communicability switched from negative in exotic species to neutral in native species (Table 1, Fig. S19). In birds, trends were qualitatively similar but the statistical support was lower. Larger-bodied native birds showed higher species-level communicability values, but the relationship was opposite and non-significant for exotic birds (Table 2, (Fig. S20)). Within local communities (Tables S1 and S2, Figs. S17 and S18), local degree was positively related to population-level communicability for plants and birds. Plant species showed the same trends as in the country-level analyses, whereas for birds, effects were also congruent but no predictors except local degree were statistically significant.

**Table 1:** Coefficients of the generalised linear model (Gamma distribution, log link) relating species-level communicability with prevalence, provenance, species degree in the metaweb, and morphological traits, for plant species. N = 102.

|  | estimate | std.error | z | p.value |
| --- | --- | --- | --- | --- |
| Intercept | -7.841 | 0.551 | -14.226 | < 0.001 |
| status (Native) | 3.001 | 0.570 | 5.264 | < 0.001 |
| fruit diameter (mm) | -8.660 | 3.250 | -2.664 | 0.008 |
| max mean height (m) | 0.204 | 0.154 | 1.326 | 0.185 |
| prevalence | 1.353 | 0.195 | 6.952 | < 0.001 |
| degree | 0.655 | 0.149 | 4.410 | < 0.001 |
| status (Native):fruit diameter (mm) | 8.622 | 3.251 | 2.652 | 0.008 |

**Table 2:** Coefficients of the generalised linear model (Tweedie distribution, log link) relating species-level communicability with prevalence, provenance, species degree in the metaweb, and body size, for bird species. N = 22.

|  | estimate | std.error | z | p.value |
| --- | --- | --- | --- | --- |
| Intercept | -3.853 | 0.837 | -4.601 | < 0.001 |
| status (Native) | 1.232 | 0.848 | 1.454 | 0.146 |
| body mass (g) | -4.077 | 2.184 | -1.867 | 0.062 |
| prevalence | 0.738 | 0.224 | 3.295 | 0.001 |
| degree | 0.881 | 0.192 | 4.581 | < 0.001 |
| status (Native):body mass (g) | 4.404 | 2.166 | 2.034 | 0.042 |

In addition to examining the statistical significance of the models, we partitioned model variances between the different predictors. We did so because our study, while based on empirical data, has a variable sample size depending on the number of grid cells in which we discretise the studied area - in such cases, p-values can potentially be subject to differing rates of type-I errors. To partition the explained variance from each analysis, current methods do not allow the inclusion of interactions between variables in generalised linear models (Lai *et al*., 2022; Lüdecke *et al*., 2020). Therefore, for this estimation we report the outcomes of models without interaction terms. This analysis confirmed that, at the local level, the most important variable was always the local population degree, both for birds (Table S4) and plants (Table S5). At the country level, species prevalence and degree in the metaweb explained the greatest variance for both groups (Tables S6 and S7), thus confirming the statistical significance trends.

### Spatial patterns of communicability

The landscape-level communicability of each grid cell (*L_G_^k^*, eq. 5) varied widely across the study area (Fig. 3). In particular, most of the territory showed comparatively low values of communicability, and only a few spatial clusters displayed clearly larger values. These spatial clusters of high communicability were nearly all in native forested areas that were surrounded by crop or pasture landscapes, as in the Whanganui National Park (the single cluster with highest communicability values, in the center-east of the North Island) or the Tararua Forest Park (southern border of the North Island).

**Fig. 3:**
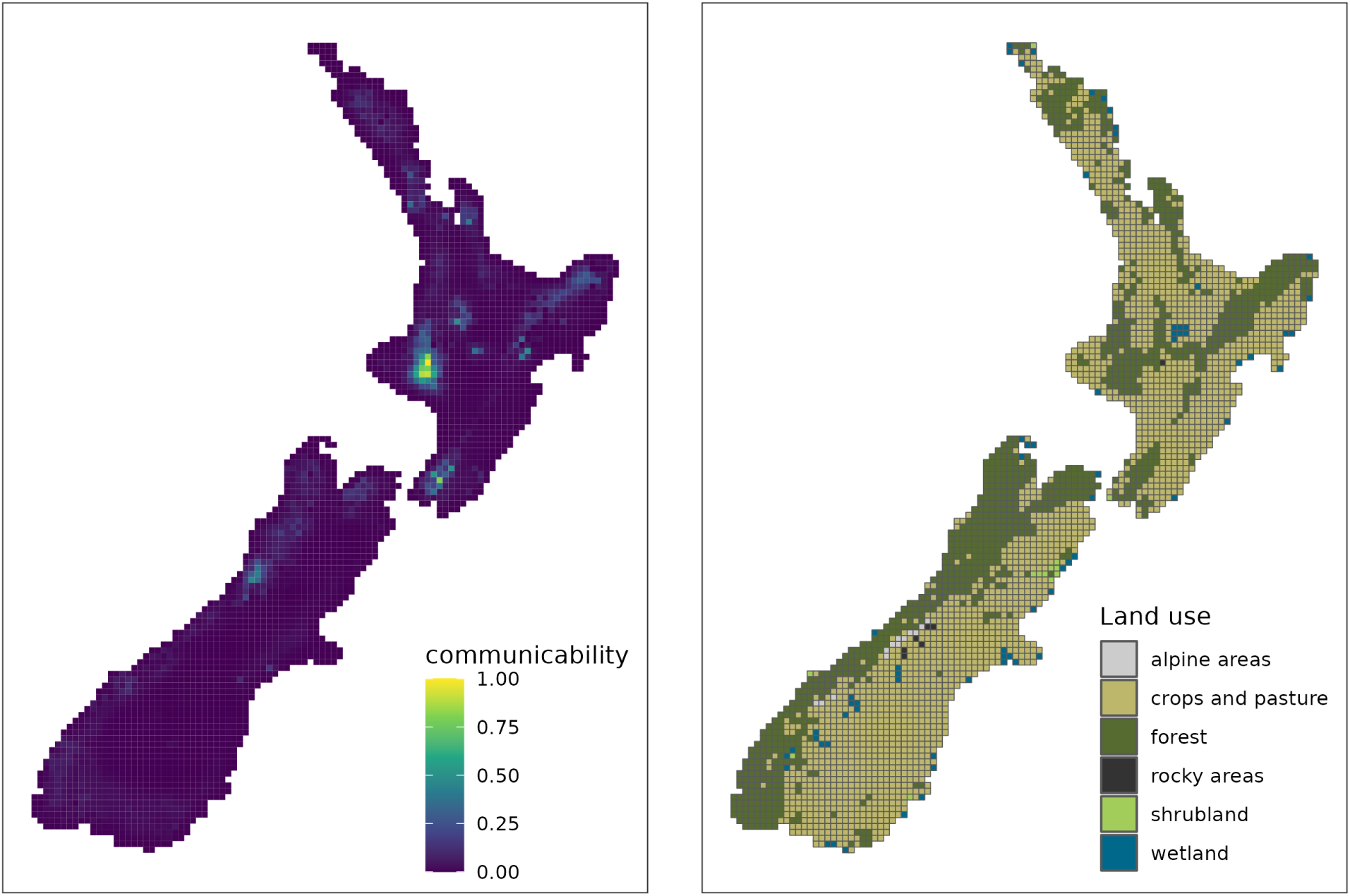
The study area (North and South Islands of Aotearoa New Zealand) showing the division in grid cells. In the left panel, aggregated communicability per cell of 10 x 10km, calculated as the sum of the communicability of each species present in the cell. In the right panel, most prevalent land use category in each cell.

Supporting this visual inspection, landscape communicability *L_G_k* was significantly and positively related to the percentage of forest and shrubland cover in the cell, and significantly but negatively related to the diversity of habitat types within the cell, measured as the Shannon index of their relative frequencies. These relationships had also a spatial signal, as there was a signficant spatial auto-correlation term in the analysis (Table 3). The influence of forest cover was likely mediated by changes in species richness, as percentage forest cover was significantly and positively correlated with the proportion of native bird species (Spearman’s *ρ* = 0.65) and with overall bird richness (*ρ* = 0.57), as well as with the proportion of native plant species (*ρ* = 0.44) and overall plant richness (*ρ* = 0.66). Shrubland cover percentage and the diversity of habitat types showed weaker correlations among them and with other variables, always being *<* 0.5 (see Fig. S16 for all pairwise correlations). We further considered the effect of forest/shrubland cover in the neighbouring area of the focal cell, as well as the diversity of habitat types in that neighbouring area, but discarded these variables due to their high correlation with their cell-level counterparts. Nevertheless, the results of a model with neighbor-only landscape variables (Table S3) showed similar patterns as our main model.

**Table 3:** Coefficients of the generalised additive model relating landscape-level communicability with landscape characteristics. The expressive degrees of freedom of the spatial smoothing term is 28.17, with an F-value of 43.21 and a p-value *<* 0.001.

|  | estimate | std.error | t | p.value |
| --- | --- | --- | --- | --- |
| Intercept | 12.657 | 0.02 | 623.008 | $< 0.001$ |
| forest cover (%) | 1.009 | 0.025 | 39.651 | $< 0.001$ |
| shrubland cover (%) | 0.407 | 0.026 | 15.733 | $< 0.001$ |
| habitat diversity | -0.198 | 0.026 | -7.773 | $< 0.001$ |

## Discussion

Quantifying the biotic effects of species and their spatial signature is essential for understanding and predicting spatial ecological dynamics within (Morris *et al*., 2005) and across habitats (Frost *et al*., 2016; Knight *et al*., 2005). A better understanding of the spatial propagation of direct and indirect effects could also provide new insights into the effects of invasive species (Vagnon *et al*., 2022), changing species distributions, the scale of land-use impacts, and disease spread in ecosystems (McCarthy *et al*., 2021; Tompkins *et al*., 2011). Here we used tools from network theory and statistical physics to show that the spatial structure of biotic interactions and population connections through dispersal can be used to approximate the spread of direct and indirect effects. More importantly, we used these tools to demonstrate the scale-dependent role of different species, traits and landscape characteristics in this tapestry of direct and indirect interactions.

We expected that a species’ potential to spread biotic effects across space should increase via multiple characteristics, including high dispersal ability, interaction generalism (i.e. high node degree), and higher abundance or spatial prevalence across the landscape. Consistent with these predictions, our analyses of a national plant-frugivore network reveals that bird and plant species that interact with many partners, within local networks and regionally, have the greatest potential to directly and indirectly affect others (i.e. highest communicability, tables 1 and 2). Moreover, and as expected, species with a widespread distribution are most important for propagating effects at large scales. These insights also align with additional theoretical results from simulated meta-communities (Supplementary Section “Communicability in simulated metacommunities”). There may exist ecologically relevant situations in which species with high communicability are not necessarily those with higher degree, e.g. in strongly disassortative networks, which may correspond to strongly nested topologies (Estrada & Hatano, 2008; Jonhson *et al*., 2013). In our system, the fact that the effect of network degree on communicability was consistent across spatial scales showed that general structural patterns of the plant-frugivore community in New Zealand were also consistent across scales, displaying a pattern of high generalism and low nestedness both locally and at larger scales when considering dispersal links. More generally, spatial scale may influence network structure in different ways (Galiana *et al*., 2018), and therefore the degree-communicability relationships across scales in other systems and interaction types remains to be explored.

Species’ traits (body size for birds, and fruit diameter for plants) were also consistent predictors of their capacity to propagate indirect effects, but this effect differed qualitatively for native and exotic species. In native birds, communicability was positively related to body size, independently of species degree (Fig. S14). This was not the case for exotic species, so we suggest that this can be a result of feeding preferences of native birds, which are known to select interaction partners more strongly on the basis of their traits (Peralta *et al*., 2020). The ecological mechanisms behind this relationship will need further investigation, but these insights show that it is possible to approximate the capacity of species to propagate ecological effects across space from a combination of morphological, functional, and possibly other traits. Further investigation of these “propagation syndromes” (Martins *et al*., 2024) in a variety of ecosystems and taxa will be important both conceptually and in an applied context, e.g. in conservation or predictions of disease spread across taxa. For example, large-bodied native species tend to have a high risk of extinction (Ripple *et al*., 2017). We have shown that such native, large-bodied species are also key in propagating ecological effects, e.g. in our study system, the Kea, *Nestor notabilis* is a locally important species (Panel A of Fig. 2) that is also highly threatened. Because indirect effects in mutualistic networks are positive in sign (in the long term), extinction of large native birds would likely have a comparatively important negative impact on the network of ecological indirect effects across the territory. Importantly, exotic bird species in Aotearoa New Zealand do not display this relationship between body size and potential propagation of indirect effects, resulting from the different local ecological roles of native and exotic species and their trait-function relationships (García *et al*., 2014; Peralta *et al*., 2020). However, exotic species are often interaction generalists (Aizen *et al*., 2008; García *et al*., 2014), a key attribute driving communicability at local and species levels. These insights thus suggest that introductions of species combining high generalism, high prevalence, and high mobility have the potential to cause far-reaching direct and indirect impacts. Further refinements of our approach may incorporate population densities when this information is available: among the insights that this could provide, we hypothesise that the higher population densities expected of large-bodied species in undisturbed relative to disturbed habitats may better disentangle the relationships between body mass, species distributions, and propagation of effects across habitats.

The analysis of communicability patterns across space (Fig. 3, Table 3) provides complementary insights to our species-level analyses by highlighting the role of community types. Areas with high forest and shrubland cover were comparatively more important for the propagation of ecological effects in space, indicating potentially widespread impacts following disturbance of forested areas. In Aotearoa New Zealand, higher forest cover is associated with a higher diversity of both plants and birds, and with a higher frequency of native bird species (Fig. S16). This suggests that communicability is partly mediated by spatial patterns of diversity, and by the differential spatial distribution of native and exotic species in our system. In other ecosystem types, communicability may thus not be necessarily related to forest cover, but to other spatial land-use configurations. The influence of the surrounding areas in our study system is qualitatively similar to that of the local community, showing that high forest/shrubland cover surrounding the focal habitat is beneficial to its diversity (Ruffell *et al*., 2017) and to its communicability. This relationship is nevertheless likely to vary across ecosystem types. For example, we would expect a more positive effect of landscape habitat diversity on spatial communicability in landscapes with a low beta diversity across different habitat types and many generalist species connecting them.

Our insights rely on the country-level spatial network of plant-frugivore interactions that we projected from systematic surveys, a distribution model extrapolating such surveys to the whole study area, and empirically-observed interactions (Fig. 1). This combined approach introduces several unavoidable sources of uncertainty in our results. First, uncertainty is measured in the goodness-of-fit of the Joint-Species Distribution Model used to infer species distributions. This model showed a very good agreement with empirical data (Supplementary Section “Spatial network validation”) and has previously shown good extrapolation ability when considering spatial structure (Norberg *et al*., 2019). Despite this potential appropriateness of the model, discrepancies in inferred distributions may impact local presences in our networks and overall prevalence. This potential uncertainty will be most relevant for rare species (Norberg *et al*., 2019), so we expect that the ordering of species according to their communicability will be robust in its first positions and may vary more strongly on its tail, where differences in communicability are quite small across species (Fig. 2). Regarding the spatial projection of interactions, our metanetwork approach assumes that two species that are known to interact will do so in every location in which they are expected to co-occur. This assumption is clearly incomplete, as the occurrence of interactions given species presences can be dependent on additional biotic or abiotic factors (Poisot *et al*., 2017). However, we lack further knowledge of what determines the realisation of plant-frugivore interactions at the spatial scales of our study, and thus our approach is a parsimonious one given the best available knowledge and data. Further, while not all potential interactions will be realized in every location, the probability of a known interaction occurring will likely increase over longer timeframes. Most importantly, in practical terms, community data with the grain and extent necessary to advance questions like those presented here is currently unavailable, and the validation performed with independent data has shown that the projected network predicts observed interactions reasonably well (Supplementary Section “Spatial network validation”), given the high degree of uncertainty associated with such data.

Notwithstanding the appropriateness of our projected network, the insights derived here must be understood as first approximations to the propagation of biotic effects across space, mediated by species interactions. The plant-frugivore network of Aotearoa New Zealand relates to demographic effects on the species considered: we assumed that effective interactions have a positive effect on the populations of the species involved (i.e. that the interactions are mutualistic), and that this effect is propagated through the subsequent plant-frugivore and bird dispersal interactions in the network. Nevertheless, the approach presented here can be used to model other effects mediated by interactions, e.g. on phenotypic selection or on behavior. The key assumption of the communicability metric we introduced is that the only driver behind the propagation of effects is the presence (or strength) of a link between a pair of species or populations, which in our case may be a plant-frugivore interaction or a dispersal link between two local populations of a given bird species. We have shown that this is a reasonable assumption for relatively simplified ecological communities, where dynamic effects are not expected to vary widely in sign or strength across species (Supplementary Section “Structural and dynamical propagation”). Another dimension that should be explored in further studies is the temporal one: variation in network topology across time or temporal delays in the propagation of effects can be potentially important modifiers of the standard communicability metric that we obtained. For example, a natural follow up would be to conduct specific observations and/or experiments about the spatiotemporal demographic effects of the species deemed to be important in analyses of effect propagation, such as here the silvereye or, locally, the kererū.

## Conclusions

Understanding how biotic effects spread in space across communities of interacting species is a fundamental unresolved question in ecology. A mechanistic understanding of effect propagation requires detailed, context-specific information on species demography, interactions, and spatial patterns, which can be overwhelmingly costly to obtain, even for moderately diverse ecological communities. We have shown that an approach based on the topology of pairwise interactions in a (meta)community allows a first approximation of such direct and indirect effects between species at varying scales. Applying this approach to a large-scale plant-frugivore network, built from existing literature and monitoring data, identified ubiquitous and generalist large-bodied species as key agents for the propagation of effects across space, and highlighted differences between native and exotic species. Whether this insight holds across ecosystem types and taxa, the temporal scales of effect propagation, or indeed, how to independently measure such indirect effects in complex natural systems, are important questions that arise from our results, and will help to advance our understanding of the spatiotemporal patterns of biodiversity.

## Methods

### Propagation of effects in ecological communities: communicability

By conceptualizing ecological communities as networks, whereby species (or populations) are nodes that are connected through links representing ecological interactions, we can analyze how community structure relates to the propagation of ecological effects. The topological structure of a network provides information on how effects on a given node can spread through the rest of the network (Wang *et al*., 2017). Any two nodes in a network can be linked through a multitude of paths, including the shortest path(s) between them and all non-shortest paths (Extended Data Fig. 4). In nodes that are not directly connected, the flow of effects is commonly analysed through their shortest path, i.e. the path that connects both nodes via the smallest number of steps. This is the case, for example, in analyses of trophic cascades along linear chains (Estes *et al*., 1998; Ford & Goheen, 2015). The length of shortest paths is captured in network metrics such as betweenness and closeness centrality. However, recent studies have emphasized the importance of indirect, non-shortest paths, in connecting nodes across different types of networks (Akbarzadeh & Estrada, 2018; García-Callejas *et al*., 2019). The combined flow across all possible paths between nodes *i* and *j* in a network is quantified by the *communicability* between them (Estrada & Hatano, 2008; Estrada *et al*., 2012). In ecological terms, it therefore may be understood as a measure of the effect of a perturbation in node *i* that reaches node *j*, and vice versa. The more connected two nodes are, via both shortest and non-shortest paths, the higher their pairwise communicability will be.

In this metric, shorter paths have larger contributions to the flow between nodes than longer ones. In particular, as defined in Estrada & Hatano (2008), if 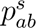 is the number of shortest paths between the nodes *a* and *b* having length *s*, and 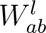 is the number of walks (i.e. sequences of nodes and links in a network) connecting *a* and *b* of length *l^ab^ > s*, the communicability between nodes *a* and *b* is defined as

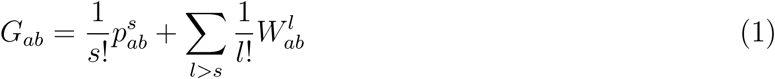

This definition implies that, as the length *l* of the paths increases, these will have an exponentially decreasing effect on the overall communicability between *a* and *b*. The formulation also allows differentiation of the effect of shortest paths and paths of arbitrary length on the metric. Further, for obtaining its aggregated value, it can be reformulated in terms of powers of the interaction matrix **A**, as

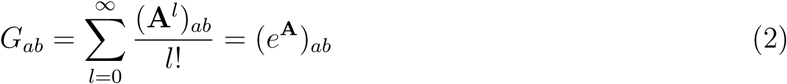

Importantly, these expressions are valid for binary matrices (Estrada & Hatano, 2008) and for positive weighted matrices (Estrada *et al*., 2012). From these pairwise values one can obtain different aggregated metrics. First, the communicability of a given node *a* in a network *k* with *R* nodes is simply the sum of pairwise values involving that node. We denote this the *population-level* communicability:

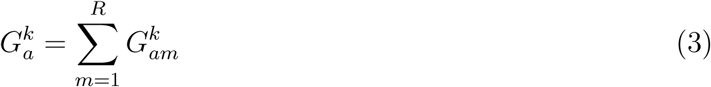

It is worth emphasising that this node- or population-level communicability is conceptually similar to centrality metrics that measure the “importance” of nodes in a network (Benzi & Klymko, 2013). However, communicability explicitly measures the effects spreading from node *i*, instead of the topological role of node *i* in the structure of the network. Furthermore, and unlike path-based centrality metrics (betweenness or closeness centrality), it accounts for all direct and indirect paths connecting two nodes instead of only shortest paths.

A further aggregated metric is available in the context of ecological networks. If nodes in a network correspond to different populations of the same species, as is potentially the case in meta-communities where different populations of a species *a* are present in different local communities (i.e. local networks) *k* (e.g. panel D of Fig. 1), an aggregated, *species-level* communicability is simply the sum of the population-level values from the different nodes:

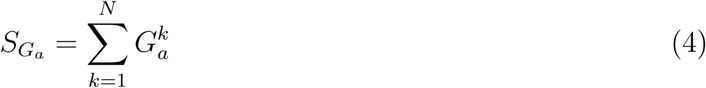

Finally, an aggregated metric at the local network level is the sum of population-level communicability values of the nodes in that local network, i.e. the total communicability of the network. We denote this the *landscape-level* communicability, as we consider each local network in a 10km grid cell an individual landscape:

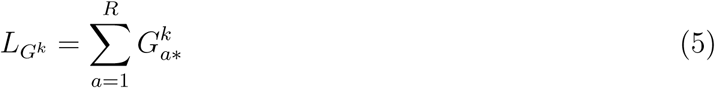

Here we focus on the applicability of these metrics to a real-world system, and in the Supplementary Section “Communicability in simulated metacommunities” we provide worked examples on obtaining and evaluating the communicability metric in *in silico* metacommunities of increasing complexity and varying structures. In addition, because communicability is a structural metric, it does not account for the dynamics of the system - e.g. whether variations in abundance across species modify indirect effects in the system, or whether negative and positive indirect effects potentially cancel each other out. Therefore, we tested the correspondence of our communicability metric with a dynamical approach to quantify net effects between species based on estimating population dynamics (Supplementary Section “Structural and dynamical propagation”).

**Fig. 4:**
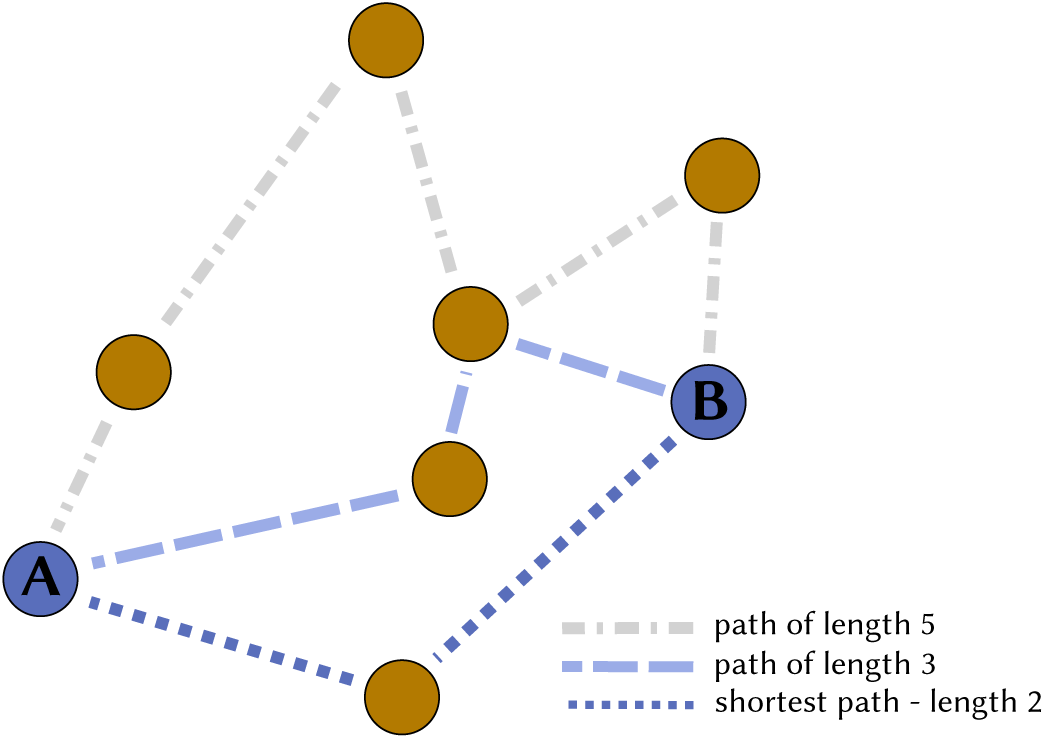
EXTENDED DATA FIGURE: Shortest path and two examples of non-shortest paths between two nodes A and B in a network. Communicability *G_AB_* approximates the effects that propagate from A to B, accounting for every path connecting them. The shortest path has the strongest influence on the metric, and every other indirect path has a contribution to communicability that decays exponentially with its path length, i.e. the number of links that are crossed. Here we draw binary links for simplicity, but the metric can also be obtained for positive-weighted networks.

### Estimating a country-level plant-frugivore network

We generated a national metaweb of plant-frugivore interactions to project a whole-country spatial interaction and dispersal network, across a grid covering the two main islands of Aotearoa New Zealand (3030 cells of 10x10 km). We then used this spatial multilayer network to evaluate how species’ potential to indirectly affect others (measured as communicability) is influenced by spatial context. Aotearoa New Zealand represents an ideal setting for such a study, because it encompasses 1) both highly fragmented landscapes and large natural areas, and 2) a large variety of habitats and climate types.

In a first step, we gathered all published plant-frugivore interactions in Aotearoa New Zealand, from the database originally obtained by Peralta *et al*. (2020). This set of potential interactions across the territory is what we term the ‘metaweb’. Detailed information on how the metaweb was built can be consulted in the original publication, but summarising, the set of potential interactions was obtained from an exhaustive literature search on bird diets, considering all birds with recorded presences in Aotearoa New Zealand. After filtering for species with available trait and geographical information, we ended up with data from 68 studies covering observations from the North (16 studies) and South (34 studies) islands as well as from smaller islands with the same species pool as the main islands (18 studies). These studies further covered all major habitat types.

To place these interactions in a spatial context, we modelled the spatial distribution of each plant and bird species using a spatially explicit Joint Species Distribution Model (Tikhonov *et al*., 2020b) fed by different sources of observational data. In particular, we gathered plant observations from datasets contained in the New Zealand National Vegetation Survey Databank (https://nvs.landcareresearch.co.nz, Wiser *et al*. (2001)). For bird species, we gathered observations from the New Zealand Department of Conservation ‘Tier 1’ monitoring program (Allen *et al*., 2013). The details of these sampling schemes are available in the references, but summarising, these datasets represent systematic observations of plant/bird individuals that we filtered to presence/absence data across the grid of the territory. We used these empirical data to project the distribution of our species over the territory by applying the joint species distribution modelling (JSDM) framework of Hierarchical Modelling of Species Communities (HMSC) (Ovaskainen & Abrego, 2020; Ovaskainen *et al*., 2017) implemented through the R-package Hmsc-R (Tikhonov *et al*., 2020b). We included in the JSDM model simultaneously the 124 species of birds and plants, and modelled their occurrences in the 3030 grid cells with occurring species. As covariates, we included the linear effects of elevation, annual precipitation, precipitation seasonality, percentage of forest cover, percentage of shrubland cover, percentage of anthropic habitat, and survey effort. To account for spatial variation in species occurrences and co-occurrences not captured by the environmental predictors, we included a spatially explicit random effect modelled through a nearest-neighbor Gaussian process (Tikhonov *et al*., 2020a). As our aim was to predict occurrence probabilities at the grid cell level, we truncated the data to presence-absence, and applied probit-regression. For grid cells with no TIER1 data, we declared the response as missing (NA) for all bird species. Similarly, for grid cells with no NVS data, we declared the response as missing (NA) for all plant species. We fitted the model with four MCMC chains, running each for 3750 iterations, out of which the first 1250 were considered as transient and the remaining thinned by 10 to yield 250 posterior samples per chain and hence 1000 posterior samples in total. We then used the fitted model to predict the probabilities for all grid cells. For these predictions, we normalized survey effort by setting it to its mean value over the grid cells, and hence our predictions correspond to the situation where the entire New Zealand would have been surveyed with constant effort. Our model showed a very good agreement with the empirical data, with AUC values *>* 0.8 for all species (see Supplementary Section “Spatial network validation”). In parallel, we obtained relevant traits for bird species from Peralta *et al*. (2020), from the AVONET database (Tobias *et al*., 2022) and body sizes from the study by Lislevand *et al*. (2007). Morphological traits for plant species were also obtained from Peralta *et al*. (2020).

Finally, we combined the metaweb of empirically-observed interactions between plants and birds from Peralta *et al*. (2020) with the spatial occurrences derived from our model to generate a spatially explicit multilayer network, including local interactions between plants and frugivores and dispersal of birds between local communities. Local interactions were assigned whenever two species that co-occurred in our model projection were known to interact in the metaweb. In other words, co-occurrence was not a sufficient criterion for inferring interactions (Blanchet *et al*., 2020); rather, co-occurring species needed to have been observed interacting in the literature. This approach assumes that all potential interactions are eventually realised locally. Yet, we acknowledge that, in empirical networks, the proportion of potential interactions that is realised can be influenced by the local environment (Laliberté & Tylianakis, 2010) and can increase with sampling intensity (Jordano, 2016). However, to our knowledge there are no consistent rules for predicting which potential interactions will be realised in a given location. Thus, we make the assumption that all possible interactions are a potential pathway for the propagation of indirect effects, including through an interaction not being realised even though both partners coexist in a site (i.e. a structural disturbance sensu Martins *et al*. (2024)). These local interactions were coded as binary. We further validated this approach by assessing how well these projected interactions match independent empirical data on 16 plant-frugivore networks across the study area. Our projected network showed a generally good agreement with empirical interactions (AUC values *>* 0.6 in 11 of the 16 empirical networks, see Supplementary Section “Spatial network validation” for details).

To connect local communities, dispersal links were assigned between two local populations of a bird species if they fell within the species’ dispersal distance. These links are of a different type to interactions, and are called ‘inter-layer’ links in multilayer network terminology (whereas interactions are ‘intra-layer’ links). We estimated bird dispersal kernels with a negative exponential function:

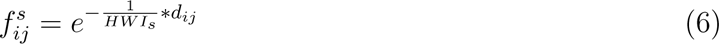

where 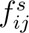 is the dispersal rate between cells *i* and *j* for species *s*, *HWI_s_* is the hand-wing index (Sheard *et al*., 2020) of species *s* and *d_ij_* is the distance between cells *i* and *j*. Two populations of a bird species (in nearby cells) were connected with a strength between [0,1] determined by their dispersal kernel and the distance between the two cells. By linking local communities in this way, we assume, for simplicity, that dispersal by birds is most important for fruiting species, and its dynamics are much faster, than dispersal of plant propagules by other means, so we ignore dispersal by plants.

### Communicability calculation

From our spatial plant-frugivore network, we obtained the population-level communicability of each local population in each cell (see details in Supplementary Section “Communicability of the Aotearoa New Zealand plant-frugivore network”) rescaled to values [0, 1]. We subsequently obtained the distribution of population-level communicability values for each species, their species-level communicability (e.g. the aggregated population-level communicability of a species across all the territory), and the landscape-level communicability (e.g. the sum of population-level communicabilities of all populations present in a given grid cell) for our spatial analyses.

### Statistical analyses

We analysed whether the communicability of a given species was related to its provenance (native vs. exotic), its network degree, and morphological traits (body mass for birds; fruit diameter and maximum mean vegetative height for plants). We selected these variables because they are ecologically meaningful, data were relatively easy to obtain compared with other traits, and were not correlated among them in either birds or plants (Figs. S14 and S15). We further explicitly tested the interaction between species provenance and body mass (birds) or fruit diameter (plants), because previous research (García *et al*., 2014; Peralta *et al*., 2020) has shown that trait matching is less important for exotic than for native species. We did not include bird gape size in our analyses, despite including fruit diameter, because it is highly correlated with body mass (Fig. S14), and our focus was not to predict interactions through trait matching, but rather analysing the importance of overarching traits associated with many life-history characteristics (Woodward *et al*., 2005). In addition, we did not include dispersal ability, quantified via the Hand-Wing Index, to avoid circularity, because that trait was used to link populations in our spatial network. We repeated these analyses for the average population communicability and the species-level communicability, to understand potential differences between local and regional drivers. For quantifying network degree, we considered the average local degree of the species, across all cells in which the species appears (including local and dispersal links). For the country-level scale, the degree metric is that of the species in the metaweb. At the regional level, we also considered species’ prevalence (the number of cells with recorded presences) as a predictor. For these analyses, we used generalised linear models with a gamma distribution and log link function, implemented in the glmmTMB package v1.1.7 (Brooks *et al*., 2017). The only exception was the GLM for bird communicability at the species level, which showed better diagnostics by fitting it with a more flexible tweedie distribution, so we kept the tweedie GLM (coefficients were very similar between gamma and tweedie GLMs, nevertheless).

We also tested whether landscape-level communicability was related to landscape characteristics: percentage of forest cover, percentage of shrubland cover, and habitat diversity (measured as the Shannon index of the relative frequency of habitat types present in the cell). We obtained spatial habitat data for New Zealand from the IUCN habitat characterisation map by Jung *et al*. (2020). We modelled these relationships with a generalised additive model with a tweedie distribution and log link function, explicitly considering spatial autocorrelation via a smoothing term on the coordinates of cell centroids. We implemented this model in the mgcv package v1.9-0. (Wood, 2011). For this and the previous models, we scaled all numerical variables before fitting and checked model consistency with the DHARMa package v0.4.6 (Hartig, 2022). All analyses were implemented in R v4.3.

## Acknowledgements

We are thankful to Hao Ran Lai, Kate Wootton, Lucas Martins, Daniel Stouffer, and Vasilis Dakos for their feedback on manuscript drafts. We acknowledge the use of data drawn from the National Vegetation Survey Databank (NVS) on 29/11/2022, and the use of data drawn from the Natural Forest plot data collected between January 2002 and March 2007 by the LUCAS programme for the Ministry for the Environment. DG-C, DP and JMT were funded by the NZ’s Biological Heritage National Science Challenge, and JMT was funded by the Ministry for Business, Innovation and Employment (programmes C09X2104 and C09X2209). OO was funded by Academy of Finland (grant no. 336212 and 345110), and the European Union: the European Research Council (ERC) under the European Union’s Horizon 2020 research and innovation programme (grant agreement No 856506; ERC-synergy project LIFEPLAN), the HORIZON-CL6-2021-BIODIV-01 project 101059492 (Biodiversity Genomics Europe), and the HORIZON-INFRA-2021-TECH-01 project 101057437 (Biodiversity Digital Twin for Advanced Modelling, Simulation and Prediction Capabilities). SL acknowledges BiodivERsA+ project RECONNECT (ANR-22-EBIP-0009-06).

## Data and code availability

The data and code used to generate the results of this study are stored in https://zenodo.org/doi/10.5281/zenodo.10812903

Development versions are stored in https://github.com/garciacallejas/propagation

## Notes

### Competing Interest Statement

The authors have declared no competing interest.

### Summary of Updates

Text updated to provide better clarification of indirect effects, of initial hypotheses, of methodologies used, and of ecological interpretation of results.

## Bibliography

Abrams, P. A., Menge, B. A., Mittelbach, G. G., Spiller, D. A. & Yodzis, P. (1996). The role of indirect effects in food webs. In: Food webs: Integration of patterns & dynamics. Springer, pp. 371–395.

Aizen, M. A., Morales, C. L. & Morales, J. M. (2008). Invasive mutualists erode native pollination webs. PLoS biology, 6, e31.

Akbarzadeh, M. & Estrada, E. (2018). Communicability geometry captures traffic flows in cities. Nature Human Behaviour, 2, 645–652.

Allen, R. B., Wright, E. F., MacLeod, C. J., Bellingham, P. J., Forsyth, D. M., Mason, N. W. H., Gormley, A. M., Marburg, A. E., MacKenzie, D. I. & McKay, M. (2013). Designing an inventory and monitoring programme for the Department of Conservation’s Natural Heritage Management System. Landcare Research Contract Report, LC1730.

Bartomeus, I., Gravel, D., Tylianakis, J. M., Aizen, M. A., Dickie, I. A. & Bernard-Verdier, M. (2016). A common framework for identifying linkage rules across different types of interactions. Functional ecology, 30, 1984–1903.

Benzi, M. & Klymko, C. (2013). Total communicability as a centrality measure. Journal of Complex Networks, 1, 124–149.

Blanchet, F. G., Cazelles, K. & Gravel, D. (2020). Co-occurrence is not evidence of ecological interactions. Ecology Letters, 23, 1050–1063.

Boesing, A. L., Nichols, E. & Metzger, J. P. (2018). Biodiversity extinction thresholds are modulated by matrix type. Ecography, 41, 1520–1533.

Brooks, M. E., Kristensen, K., van Benthem, K. J., Magnusson, A., Berg, C. W., Nielsen, A., Skaug, H. J., Maechler, M. & Bolker, B. M. (2017). glmmTMB balances speed and flexibility among packages for zero-inflated generalized linear mixed modeling. The R Journal, 9, 378–400.

Cosmo, L. G., Assis, A. P. A., de Aguiar, M. A. M., Pires, M. M., Valido, A., Jordano, P., Thompson, J. N., Bascompte, J. & Guimarães, P. R. (2023). Indirect effects shape species fitness in coevolved mutualistic networks. Nature, in press.

Donohue, I., Petchey, O. L., Kéfi, S., Génin, A., Jackson, A. L., Yang, Q. & O’Connor, N. E. (2017). Loss of predator species, not intermediate consumers, triggers rapid and dramatic extinction cascades. Global Change Biology, 23, 2962–2972.

Estes, J. A., Tinker, M. T., Williams, T. M. & Doak, D. F. (1998). Killer Whale Predation on Sea Otters Linking Oceanic and Nearshore Ecosystems. Science, 282, 473–476.

Estrada, E. & Hatano, N. (2008). Communicability in complex networks. Physical Review E, 77, 036111.

Estrada, E., Hatano, N. & Benzi, M. (2012). The physics of communicability in complex networks. Physics Reports, 514, 89–119.

Ford, A. T. & Goheen, J. R. (2015). Trophic Cascades by Large Carnivores: A Case for Strong Inference and Mechanism. Trends in Ecology & Evolution, 30, 725–735.

Frost, C. M., Didham, R. K., Rand, T. A., Peralta, G. & Tylianakis, J. M. (2015). Community-level net spillover of natural enemies from managed to natural forest. Ecology, 96, 193–202.

Frost, C. M., Peralta, G., Rand, T. A., Didham, R. K., Varsani, A. & Tylianakis, J. M. (2016). Apparent competition drives community-wide parasitism rates and changes in host abundance across ecosystem boundaries. Nature Communications, 7, 12644.

Galiana, N., Lurgi, M., Claramunt-López, B., Fortin, M.-J., Leroux, S., Cazelles, K., Gravel, D. & Montoya, J. M. (2018). The spatial scaling of species interaction networks. Nature Ecology & Evolution, 2, 782–790.

García, D., Martínez, D., Stouffer, D. B. & Tylianakis, J. M. (2014). Exotic birds increase generalization and compensate for native bird decline in plant–frugivore assemblages. Journal of Animal Ecology, 83, 1441–1450.

García-Callejas, D., Molowny-Horas, R., Araújo, M. B. & Gravel, D. (2019). Spatial trophic cascades in communities connected by dispersal and foraging. Ecology, 100, e02820.

Gounand, I., Harvey, E., Little, C. J. & Altermatt, F. (2018). Meta-Ecosystems 2.0: Rooting the Theory into the Field. Trends in Ecology & Evolution, 33, 36–46.

Gravel, D., Massol, F. & Leibold, M. A. (2016). Stability and complexity in model meta-ecosystems. Nature communications, 7, 12457.

Guimarães Jr, P. R., Pires, M. M., Jordano, P., Bascompte, J. & Thompson, J. N. (2017). Indirect effects drive coevolution in mutualistic networks. Nature, 550, 511–514.

Hackett, T. D., Sauve, A. M. C., Davies, N., Montoya, D., Tylianakis, J. M. & Memmott, J. (2019). Reshaping our understanding of species’ roles in landscape-scale networks. Ecology Letters, 22, 1367–1377.

Hartig, F. (2022). DHARMa: Residual Diagnostics for Hierarchical (Multi-Level / Mixed) Regression Models. URL https://CRAN.R-project.org/package=DHARMa. R package version 0.4.6.

Harvey, E., Gounand, I., Ganesanandamoorthy, P. & Altermatt, F. (2016). Spatially cascading effect of perturbations in experimental meta-ecosystems. Proceedings of the Royal Society B: Biological Sciences, 283, 20161496.

Higashi, M. & Nakajima, H. (1995). Indirect effects in ecological interaction networks i. the chain rule approach. Mathematical Biosciences, 130, 99–128.

Holt, R. D. & Bonsall, M. B. (2017). Apparent competition. Annual Review of Ecology, Evolution, and Systematics, 48, 447–471.

Jacobson, B. & Peres-Neto, P. R. (2010). Quantifying and disentangling dispersal in metacommunities: How close have we come? how far is there to go? Landscape Ecology, 25, 495–507.

Jonhson, S., Domínguez-García, V. & Muñoz, M. A. (2013). Factors determining nestedness in complex networks. PLOS ONE, 8, e74025.

Jordano, P. (2016). Sampling networks of ecological interactions. Functional Ecology, 30, 1883– 1893.

Jung, M., Dahal, P. R., Butchart, S. H., Donald, P. F., De Lamo, X., Lesiv, M., Kapos, V., Rondinini, C. & Visconti, P. (2020). A global map of terrestrial habitat types. Scientific data, 7, 256.

Knight, T. M., McCoy, M. W., Chase, J. M., McCoy, K. A. & Holt, R. D. (2005). Trophic cascades across ecosystems. Nature, 437, 880–883.

Kortsch, S., Primicerio, R., Fossheim, M., Dolgov, A. V. & Aschan, M. (2015). Climate change alters the structure of arctic marine food webs due to poleward shifts of boreal generalists. Proceedings of the Royal Society B: Biological Sciences, 282, 20151546.

Lai, J., Zou, Y., Zhang, S., Zhang, X. & Mao, L. (2022). glmm.hp: an r package for computing individual effect of predictors in generalized linear mixed models. Journal of Plant Ecology, 15, 1302–1307.

Laliberté, E. & Tylianakis, J. M. (2010). Deforestation homogenizes tropical parasitoid–host networks. Ecology, 91, 1740–1747.

Lislevand, T., Figuerola, J. & Sźekely, T. (2007). Avian Body Sizes in Relation to Fecundity, Mating System, Display Behavior, and Resource Sharing. Ecology, 88, 1605–1605.

Lüdecke, D., Ben-Shachar, M. S., Patil, I. & Makowski, D. (2020). Extracting, computing and exploring the parameters of statistical models using R. Journal of Open Source Software, 5, 2445.

Mariani, M. S., Ren, Z.-M., Bascompte, J. & Tessone, C. J. (2019). Nestedness in complex networks: Observation, emergence, and implications. Physics Reports, 813, 1–90.

Martins, L. P., García-Callejas, D., Lai, H. R., Wootton, K. L. & Tylianakis, J. M. (2024). The propagation of disturbances in ecological networks. Trends in Ecology and Evolution, 39, 558– 570.

Marvier, M., Kareiva, P. & Neubert, M. G. (2004). Habitat destruction, fragmentation, and disturbance promote invasion by habitat generalists in a multispecies metapopulation. Risk Analysis: An International Journal, 24, 869–878.

McCann, K. S., Cazelles, K., MacDougall, A. S., Fussmann, G. F., Bieg, C., Cristescu, M., Fryxell, J. M., Gellner, G., Lapointe, B. & Gonzalez, A. (2021). Landscape modification and nutrientdriven instability at a distance. Ecology Letters, 24, 398–414.

McCann, K. S., Rasmussen, J. B. & Umbanhowar, J. (2005). The dynamics of spatially coupled food webs. Ecology Letters, 8, 513–523.

McCarthy, J. K., Wiser, S. K., Bellingham, P. J., Beresford, R. M., Campbell, R. E., Turner, R. & Richardson, S. J. (2021). Using spatial models to identify refugia and guide restoration in response to an invasive plant pathogen. Journal of Applied Ecology, 58, 192–201.

McCoy, M. W., Barfield, M. & Holt, R. D. (2009). Predator shadows: Complex life histories as generators of spatially patterned indirect interactions across ecosystems. Oikos, 118, 87–100.

Montoya, J. M., Woodward, G., Emmerson, M. C. & Soĺe, R. V. (2009). Press Perturbations and Indirect Effects in Real Food Webs. Ecology, 90, 2426–2433.

Morris, R. J., Lewis, O. T. & Godfray, H. C. J. (2004). Experimental evidence for apparent competition in a tropical forest food web. Nature, 428, 310–313.

Morris, R. J., Lewis, O. T. & Godfray, H. C. J. (2005). Apparent competition and insect community structure: Towards a spatial perspective. Annales Zoologici Fennici, 42, 449–462.

Nakajima, H. & Higashi, M. (1995). Indirect effects in ecological interaction networks ii. the conjugate variable approach. Mathematical Biosciences, 130, 129–150.

Norberg, A., Abrego, N., Blanchet, F. G., Adler, F. R., Anderson, B. J., Anttila, J., Araújo, M. B., Dallas, T., Dunson, D., Elith, J., Foster, S. D., Fox, R., Franklin, J., Godsoe, W., Guisan, A., O’Hara, B., Hill, N. A., Holt, R. D., Hui, F. K. C., Husby, M., Kålås, J. A., Lehikoinen, A., Luoto, M., Mod, H. K., Newell, G., Renner, I., Roslin, T., Soininen, J., Thuiller, W., Vanhatalo, J., Warton, D., White, M., Zimmermann, N. E., Gravel, D. & Ovaskainen, O. (2019). A comprehensive evaluation of predictive performance of 33 species distribution models at species and community levels. Ecological Monographs, 89, e01370.

Novak, M., Yeakel, J. D., Noble, A. E., Doak, D. F., Emmerson, M., Estes, J. A., Jacob, U., Tinker, M. T. & Wootton, J. T. (2016). Characterizing species interactions to understand press perturbations: What is the community matrix? Annual Review of Ecology, Evolution, and Systematics, 47, 409–432.

Ovaskainen, O. & Abrego, N. (2020). Joint species distribution modelling: With applications in R. Cambridge University Press.

Ovaskainen, O., Tikhonov, G., Norberg, A., Guillaume Blanchet, F., Duan, L., Dunson, D., Roslin, T. & Abrego, N. (2017). How to make more out of community data? A conceptual framework and its implementation as models and software. Ecology Letters, 20, 561–576.

Peralta, G., Perry, G. L. W., Vázquez, D. P., Dehling, D. M. & Tylianakis, J. M. (2020). Strength of niche processes for species interactions is lower for generalists and exotic species. Journal of Animal Ecology, 89, 2145–2155.

Pillai, P., Gonzalez, A. & Loreau, M. (2011). Metacommunity theory explains the emergence of food web complexity. Proceedings of the National Academy of Sciences, 108, 19293–19298.

Pires, M. M., O’Donnell, J. L., Burkle, L. A., Díaz-Castelazo, C., Hembry, D. H., Yeakel, J. D., Newman, E. A., Medeiros, L. P., de Aguiar, M. A. M. & Guimarães, P. R. (2020). The indirect paths to cascading effects of extinctions in mutualistic networks. Ecology, 101, e03080.

Poisot, T., Gueveneux-Julien, C., Fortin, M.-J., Gravel, D. & Legendre, P. (2017). Hosts, parasites, and their interactions respond to different climatic variables. Global Ecology and Biogeography, 26, 942–951.

Raffard, A., Bestion, E., Cote, J., Haegeman, B., Schtickzelle, N. & Jacob, S. (2022). Dispersal syndromes can link intraspecific trait variability and meta-ecosystem functioning. Trends in Ecology & Evolution, 37, 322–331.

Ripple, W. J., Estes, J. A., Schmitz, O. J., Constant, V., Kaylor, M. J., Lenz, A., Motley, J. L., Self, K. E., Taylor, D. S. & Wolf, C. (2016). What is a Trophic Cascade? Trends in Ecology & Evolution, 31, 842–849.

Ripple, W. J., Wolf, C., Newsome, T. M., Hoffmann, M., Wirsing, A. J. & McCauley, D. J. (2017). Extinction risk is most acute for the world’s largest and smallest vertebrates. Proceedings of the National Academy of Sciences, 114, 10678–10683.

Ruffell, J., Clout, M. N. & Didham, R. K. (2017). The matrix matters, but how should we manage it? estimating the amount of high-quality matrix required to maintain biodiversity in fragmented landscapes. Ecography, 40, 171–178.

Sheard, C., Neate-Clegg, M. H. C., Alioravainen, N., Jones, S. E. I., Vincent, C., MacGregor, H. E. A., Bregman, T. P., Claramunt, S. & Tobias, J. A. (2020). Ecological drivers of global gradients in avian dispersal inferred from wing morphology. Nature Communications, 11, 2463.

Subalusky, A. L. & Post, D. M. (2019). Context dependency of animal resource subsidies. Biological reviews, 94, 517–538.

Tack, A. J. M., Gripenberg, S. & Roslin, T. (2011). Can we predict indirect interactions from quantitative food webs? – an experimental approach. Journal of Animal Ecology, 80, 108–118.

Tikhonov, G., Duan, L., Abrego, N., Newell, G., White, M., Dunson, D. & Ovaskainen, O. (2020a). Computationally efficient joint species distribution modeling of big spatial data. Ecology, 101, e02929.

Tikhonov, G., Opedal, Ø. H., Abrego, N., Lehikoinen, A., de Jonge, M. M., Oksanen, J. & Ovaskainen, O. (2020b). Joint species distribution modelling with the r-package hmsc. Methods in ecology and evolution, 11, 442–447.

Tobias, J. A., Sheard, C., Pigot, A. L., Devenish, A. J. M., Yang, J., Sayol, F., Neate-Clegg, M. H. C., Alioravainen, N., Weeks, T. L., Barber, R. A., Walkden, P. A., MacGregor, H. E. A., Jones, S. E. I., Vincent, C., Phillips, A. G., Marples, N. M., Montaño-Centellas, F. A., Leandro-Silva, V., Claramunt, S., Darski, B., Freeman, B. G., Bregman, T. P., Cooney, C. R., Hughes, E. C., Capp, E. J. R., Varley, Z. K., Friedman, N. R., Korntheuer, H., Corrales-Vargas, A., Trisos, C. H., Weeks, B. C., Hanz, D. M., Töpfer, T., Bravo, G. A., Remeš, V., Nowak, L., Carneiro, L. S., Moncada R., A. J., Matysioková, B., Baldassarre, D. T., Martínez-Salinas, A., Wolfe, J. D., Chapman, P. M., Daly, B. G., Sorensen, M. C., Neu, A., Ford, M. A., Mayhew, R. J., Fabio Silveira, L., Kelly, D. J., Annorbah, N. N. D., Pollock, H. S., Grabowska-Zhang, A. M., McEntee, J. P., Carlos T. Gonzalez, J., Meneses, C. G., Muñoz, M. C., Powell, L. L., Jamie, G. A., Matthews, T. J., Johnson, O., Brito, G. R. R., Zyskowski, K., Crates, R., Harvey, M. G., Jurado Zevallos, M., Hosner, P. A., Bradfer-Lawrence, T., Maley, J. M., Stiles, F. G., Lima, H. S., Provost, K. L., Chibesa, M., Mashao, M., Howard, J. T., Mlamba, E., Chua, M. A. H., Li, B., Gómez, M. I., García, N. C., Päckert, M., Fuchs, J., Ali, J. R., Derryberry, E. P., Carlson, M. L., Urriza, R. C., Brzeski, K. E., Prawiradilaga, D. M., Rayner, M. J., Miller, E. T., Bowie, R. C. K., Lafontaine, R.-M., Scofield, R. P., Lou, Y., Somarathna, L., Lepage, D., Illif, M., Neuschulz, E. L., Templin, M., Dehling, D. M., Cooper, J. C., Pauwels, O. S. G., Analuddin, K., Fjeldså, J., Seddon, N., Sweet, P. R., DeClerck, F. A. J., Naka, L. N., Brawn, J. D., Aleixo, A., Böhning-Gaese, K., Rahbek, C., Fritz, S. A., Thomas, G. H. & Schleuning, M. (2022). AVONET: Morphological, ecological and geographical data for all birds. Ecology Letters, 25, 581–597.

Tompkins, D. M., Dunn, A. M., Smith, M. J. & Telfer, S. (2011). Wildlife diseases: from individuals to ecosystems. Journal of Animal Ecology, 80, 19–38.

Vagnon, C., Rohr, R. P., Bersier, L.-F., Cattańeo, F., Guillard, J. & Frossard, V. (2022). Combining food web theory and population dynamics to assess the impact of invasive species. Frontiers in Ecology and Evolution, 10, 913954.

Wang, W., Tang, M., Stanley, H. E. & Braunstein, L. A. (2017). Unification of theoretical approaches for epidemic spreading on complex networks. Reports on progress in physics, 80, 036603.

Wiser, S. K., Bellingham, P. J. & Burrows, L. E. (2001). Managing biodiversity information: Development of New Zealand’s National Vegetation Survey databank. New Zealand Journal of Ecology, 25, 1–17.

Wood, S. N. (2011). Fast stable restricted maximum likelihood and marginal likelihood estimation of semiparametric generalized linear models. Journal of the Royal Statistical Society (B), 73, 3–36.

Woodward, G., Ebenman, B., Emmerson, M., Montoya, J. M., Olesen, J. M., Valido, A. & Warren, P. H. (2005). Body size in ecological networks. Trends in Ecology and Evolution, 20, 402–409.

Wootton, J. T. (2002). Indirect effects in complex ecosystems: Recent progress and future challenges. Journal of Sea Research, 48, 157–172.

Wootton, K. L., Curtsdotter, A., Bommarco, R., Roslin, T. & Jonsson, T. (2023). Food webs coupled in space: Consumer foraging movement affects both stocks and fluxes. Ecology, 104, e4101.

Wotton, D. M. & Kelly, D. (2012). Do larger frugivores move seeds further? body size, seed dispersal distance, and a case study of a large, sedentary pigeon. Journal of Biogeography, 39, 1973–1983.

