## Supplementary material for "Species traits and landscape structure can drive scale-dependent propagation of effects in ecosystems"

**Supplementary Section: Communicability in simulated metacommunities** **2**

**Supplementary Section: Structural and dynamical propagation** **9**

**Supplementary Section: Spatial network validation** **16**

**Supplementary Section: Communicability of the Aotearoa New Zealand plant-frugivore network** **20**

**Supplementary Section: Correlation between species-level variables** **22**

**Supplementary Section: Correlation between landscape variables** **24**

**Supplementary Section: Supplementary regressions** **25**

**Supplementary Section: Variable importance estimation** **26**

**Supplementary Figures** **28**

### Supplementary Section: Communicability in simulated meta-communities

To test how species traits and landscape structure influence the propagation of effects in a controlled setting, we simulated gridded landscapes populated by virtual species, with varying levels of three key parameters: the spatial distribution of species, the distribution of interactions between species, and their dispersal abilities. The first step was to generate the virtual landscapes, which we model as meta-communities in a multi-layer network framework. Within the multi-layer network, nodes are species and there are two types of links: interactions within a grid cell and dispersal among grid cells (with dispersal links connecting spatially separated populations of the same species).

We created grids of 100 cells ($10 \times 10$) and imputed a randomly generated value in the interval [0,1] to each cell, representing a single environmental dimension. To test the influence of landscape configuration, we generated landscapes with varying levels of spatial autocorrelation $c \in [0.1, 0.9]$ in their environmental dimension (Fig. S1), analogous to a habitat fragmentation gradient. In parallel to creating these gridded landscapes, we generated regional pools of 30 species, where each species was randomly assigned three uncorrelated values. First, a numeric value $n \in [0, 1]$ representing its optimum in the landscape environmental dimension defined above. Second, another numerical value to each species representing its dispersal potential. The dispersal potential for each species was sampled from an exponential distribution, whose rate parameter $d \in [0.1, 0.9]$ is the control parameter of the gradient in dispersal abilities (Fig. S2). Third, a measure of the species' degree (i.e. its number of interaction partners or links) in the network of interactions among the 30 species. Specifically, we obtained species' degrees by sampling from poisson distributions with means in the interval $\lambda \in [3, 15]$. The metawebs of species interactions (the networks of all potential interactions between the species present in the meta-community) were thus built from these species degree sequences by assigning links randomly among species (Fig. S3).

For each simulated lanscape, we finally placed the virtual species in grid cells according to their environmental optimum, whereby species $i$, with environmental optimum $e_i$, would be present in all cells with values $e_i \pm 0.2$. Two species were linked (i.e. interacted) in a cell if they did so in the

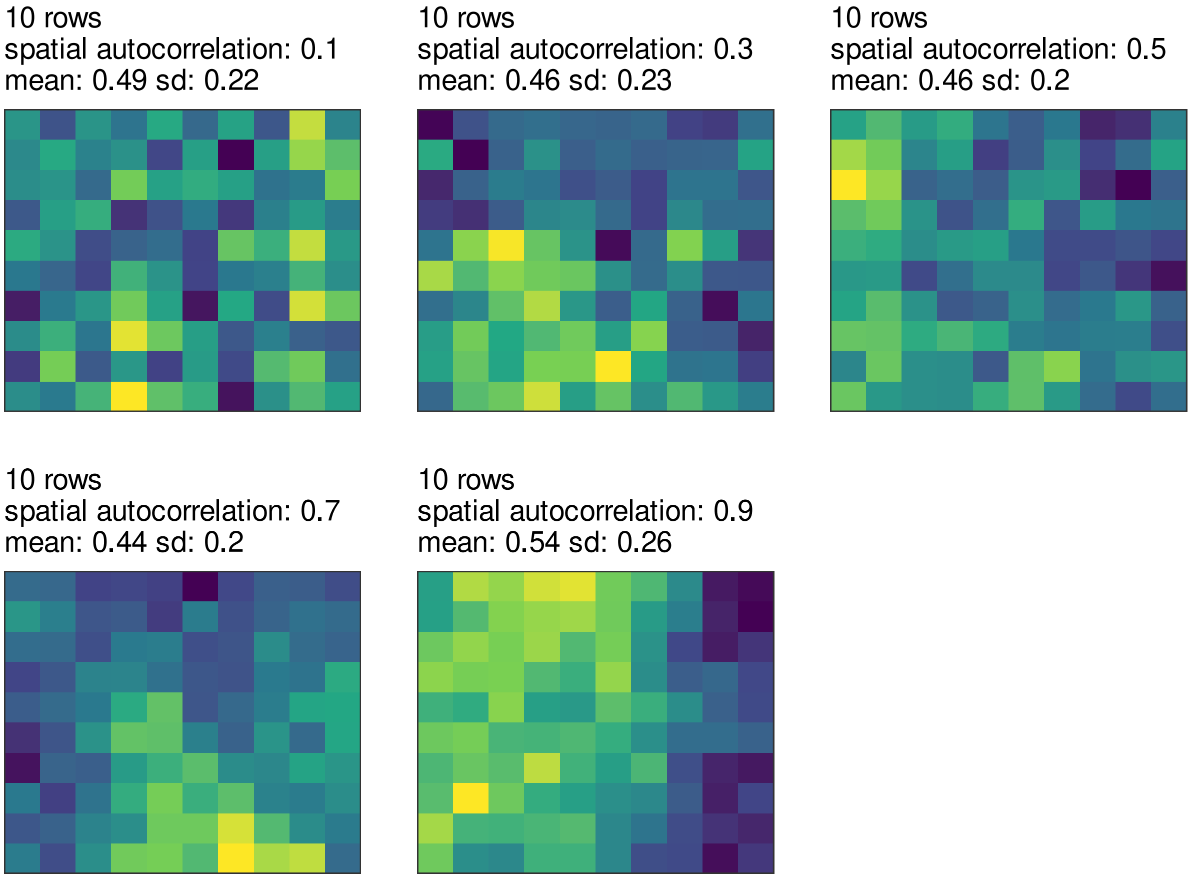


Fig. S1: Virtual landscapes with 100 cells and a single environmental dimension. Each panel shows a realization with a given level of spatial autocorrelation in the environmental dimension. The distribution of the environmental dimension values across the landscapes is maintained in the different levels of spatial autocorrelation.

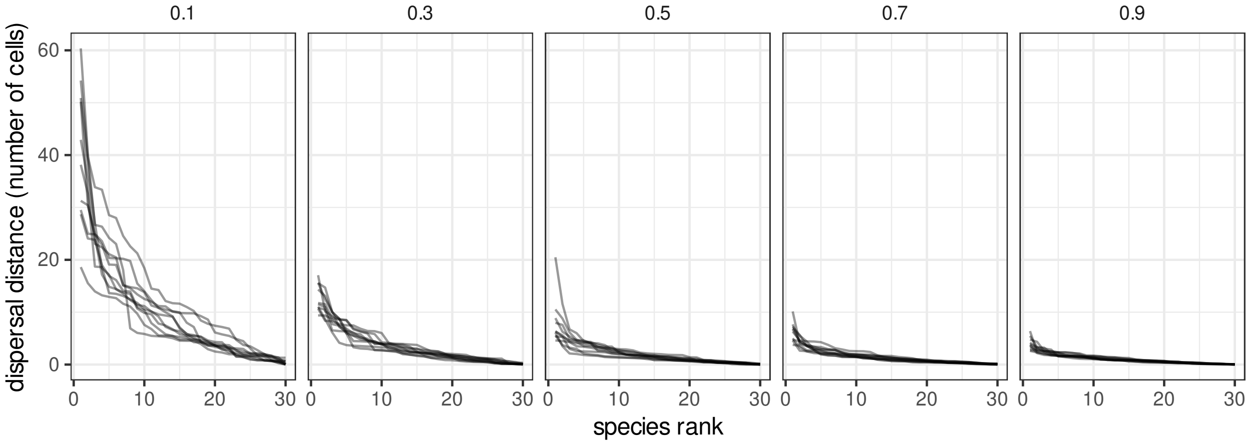


Fig. S2: Dispersal distances of species simulated under different negative exponental distributions, with rate parameters between 0.1 and 0.9. Each line in each panel is a simulated set of 30 species.

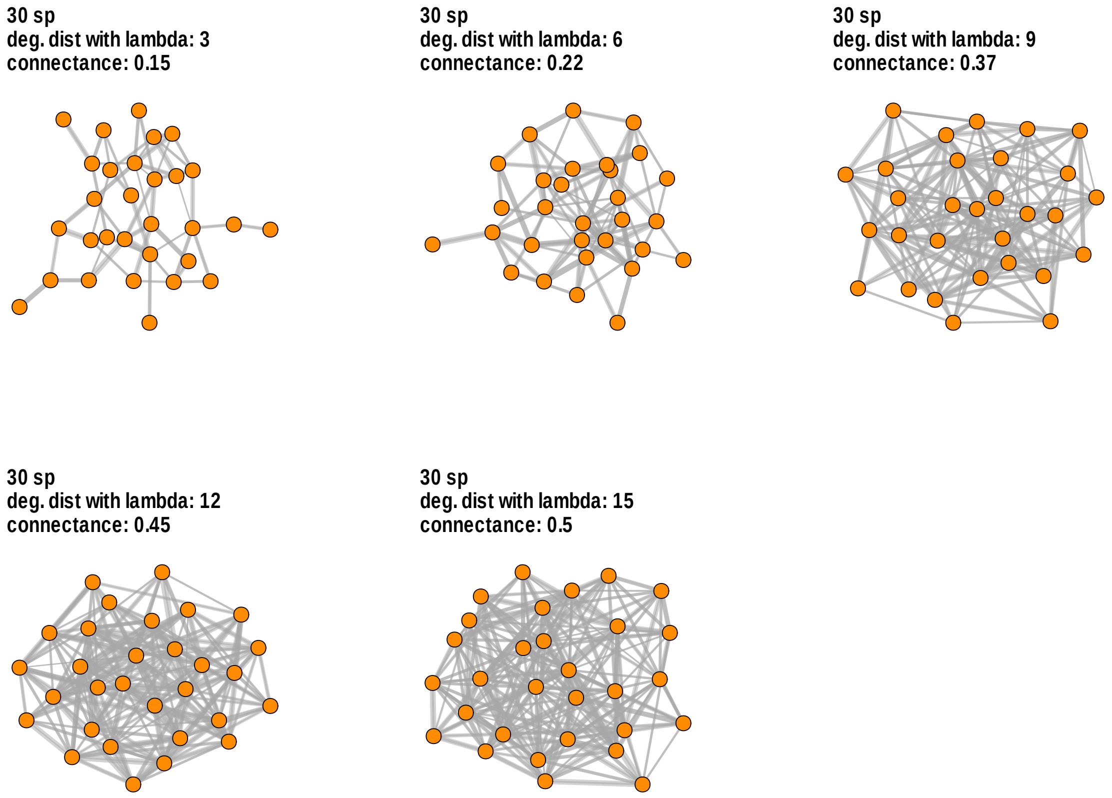


Fig. S3: Simulated metawebs of 30 species. Each panel shows a realization with a given degree distribution, where the degree distribution is taken from a poisson distribution with mean equal to lambda.

metaweb, such that each cell in the landscape had a local interaction network that was a subset of the metaweb. Finally, the dispersal links between occurrences of a given species in different cells were imputed according to its dispersal potential: if a population of a species was present in a cell that was within the dispersal range of another population of that same species in another cell, a dispersal link was established between both populations. For simplicity, we did not include spatial links connecting different species in our networks. In our simulated landscapes, interaction and dispersal links are both binary, so communicability is equally affected by dispersal and interaction links. Thus, each simulated landscape resulted from a combination of 1) spatial autocorrelation in an environmental dimension ($c$), which influenced the distribution of the virtual species, 2) connectivity of the corresponding metaweb ($\lambda$), and 3) distribution of dispersal distances ($d$).

Note that this approach is static, i.e. we do not model occupancy dynamics, but rather each virtual landscape represents a single snapshot of a metacommunity given by these three control parameters. We generated a gradient of 5 values for each of these parameters, for a total of $5^3 = 125$ combinations. For each of these combinations we generated 10 landscape replicates, for a total of 1250 simulated landscapes. Lastly, for each of these, we obtained the species-level aggregated communicability for each species $a$, $S_{G_a}$ (Eq. 4 of main text), which informs about the contribution of species $a$ in propagating effects across the landscape, and the (normalised) landscape-level communicability $L_{G^k}$ (Eq. 5 of main text).

We assessed how landscape-level communicability $L_{G^k}$ was influenced by the three control parameters of our simulated landscapes, namely 1) landscape configuration (i.e. spatial autocorrelation of the environmental dimension), 2) metaweb structure summarised via species degree and its corresponding distribution, which implies overall metaweb connectance (Poisot & Gravel, 2014), and 3) dispersal capacity in the species pool. Likewise, at the species level, we evaluated the relationship between species' communicability $S_{G_a}$ and three species-level properties: 1) prevalence (the number of occuppied cells in the simulated landscape), 2) generality (degree in the metaweb), and 3) dispersal distance. Since these results are obtained from *in silico* observations, we analysed them qualitatively rather than performing statistical tests on the simulation outputs (White *et al.*, 2014).

The results show that communicability was mainly influenced by the structure of the metaweb (Fig. S4): higher species mean degree (i.e. species being interaction generalists), with corresponding higher metaweb connectance, resulted in higher communicability in the simulated landscapes. There was a smaller but positive signal of increasing dispersal distance, and lastly, landscape configuration (autocorrelation of the environmental dimension) showed no effect on the communicability values.

At the species level, communicability was only influenced by a species' prevalence (the number of presences in the landscape), with an average Spearman's correlation coefficient of 0.73 (Fig. S5). Species' generality (degree in the metaweb) and dispersal distance were mostly uncorrelated with their aggregated communicability. Together, these simulations indicate that a species' prevalence in the landscape has the greatest impact on its ability to indirectly affect others, whereas its generalism and dispersal ability are more important at the community scale (i.e. when many species change in these traits).

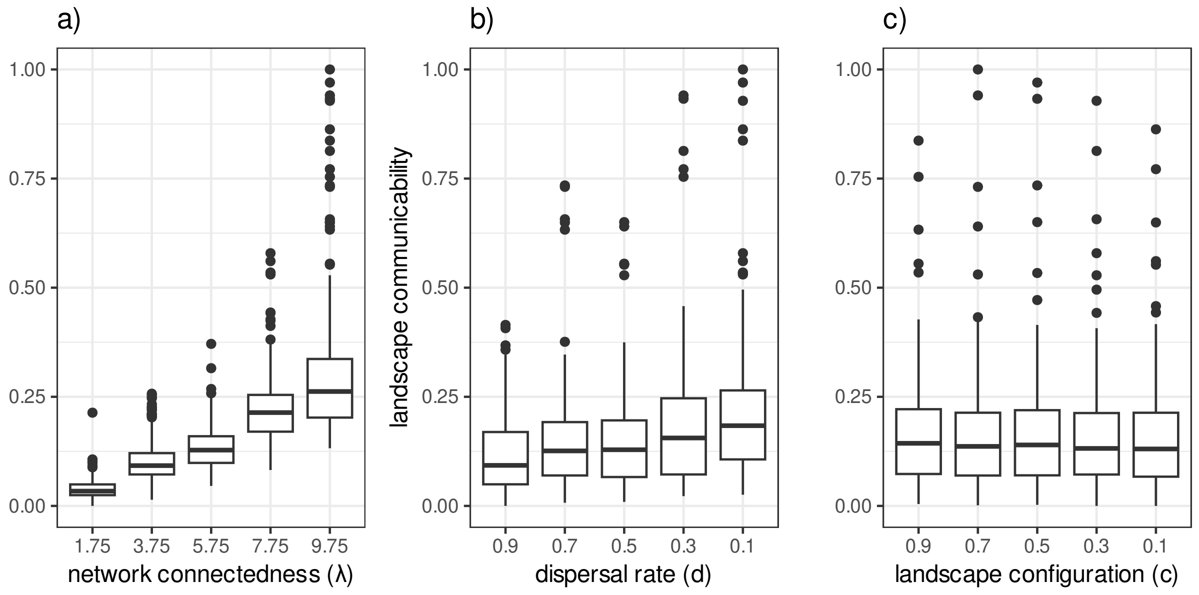


Fig. S4: Network-level communicability in simulated landscapes and its variation with three landscape-level factors: a) connectedness of species interactions ($\lambda$ of the poisson distribution of links per species, i.e. degree, in the metanetwork); b) dispersal ability of species (rate parameter of an exponential distribution, with higher values indicating that species disperse greater distances); c) landscape configuration (spatial autocorrelation in the suitability of the environment for different species). In the boxplots, the horizontal black lines represent the median, the lower and upper hinges correspond to the 25th and 75th percentiles, and the vertical lines extend to the largest/smallest value up to 1.5 times the interquartile range (distance between 25th and 75th percentiles). N = 250 in each boxplot.

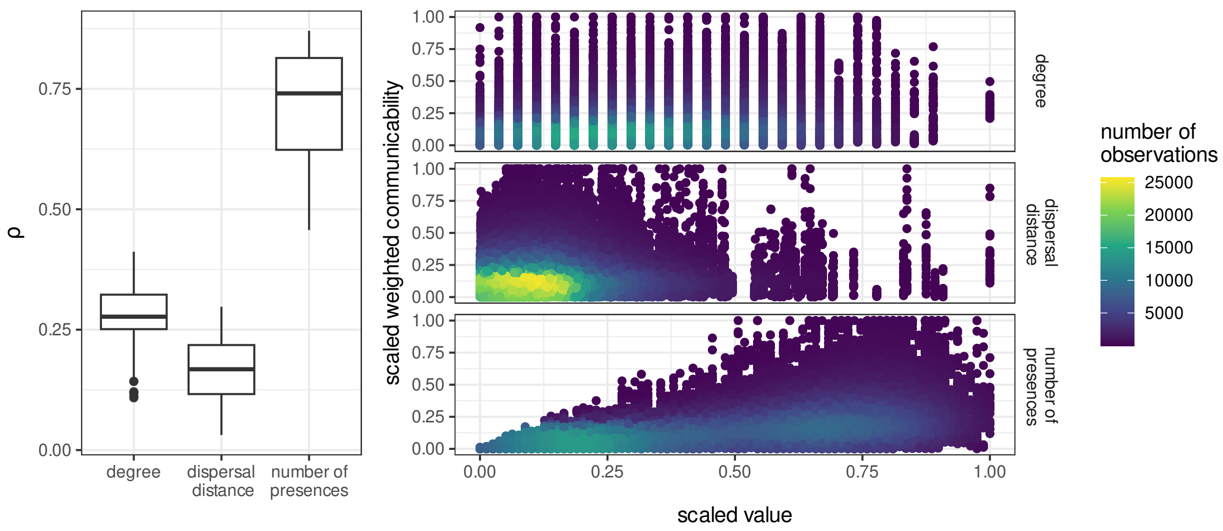


Fig. S5: Species-level communicability and its correlation with species traits. The left panel shows the distribution of Spearman's $\rho$ correlation values between communicability and species' generality (degree), dispersal rate, and prevalence (the number of presences in the virtual landscape), respectively. The right panel shows how observations are distributed across these factors. N = 125 in boxplots of left panel, 34745 in each scatterplot of right panel.

### Supplementary Section: Structural and dynamical propagation

Communicability is a structural property of networks, i.e. it does not account for the dynamics of the nodes (populations, in our conceptualization), and how these dynamics interact to amplify or dampen indirect effects throughout an ecological community.

In community ecology, in particular in the food web literature, it is well established that population dynamics can generate cascades of indirect effects driven by the sign and strength of interactions between the different species in the network. However, these estimates of indirect effects require a considerable amount of empirical data when estimated in empirical communities, as well as strong assumptions about the dynamics of the system. In particular, projecting the dynamics of the system with mathematical models requires, at the very least, estimates of intrinsic growth rates for all species in the community, as well as the sign and per capita strength of every pairwise direct interaction, which are assumed to be constant. If the system is spatially explicit, estimates of dispersal coefficients between patches are also required (e.g. García-Callejas *et al.* (2019)). From the numerical estimation of the steady-state solutions of the system, the Jacobian matrix informs about the direct effect of the average species $j$ individual on the population growth rate of species $i$. Likewise, the negative of the inverse Jacobian matrix of the system at equilibrium informs of the net effects, including both direct and indirect ones, of a sustained unit increase in population growth rate of species $j$ on the abundance of species $i$ (Novak *et al.*, 2016).

It is unknown whether estimates of indirect effects between pairs of species, estimated via the negative inverse Jacobian matrix of the dynamical system, are correlated to structural estimates of effects propagation such as the communicability metric used in this study. In this section we evaluate this relationship by comparing the communicability and the dynamical net effects across species pairs in simulated communities of varying richness.

We simulated the dynamics of three types of communities: competitive, mutualistic, and food webs with varying levels of species richness, and numerically estimated the net effects between each pair of species as well as their pairwise communicability. For each type of community and level of richness (20, 40, 60, 80, or 100 species), we generated 100 networks, for a total of $3 * 5 * 100 = 1500$ networks. Here we specify the generative model and simulations for the different communities, and discuss the results obtained.

### Community generative models

The overall workflow is similar in all cases: for each community simulated, first we generated an interaction matrix $A$ according to a given set of rules, then we assigned interaction strenghts and other model parameters (e.g. growth rates, initial abundances), and finally we solved the associated dynamical system to obtain pairwise net effects. For all models, the values of the associated parameters and the full R code to reproduce the results are available at
https://github.com/garciacallejas/propagation,
in particular in the script `R/sim_11_supp_mat_test_communicability_dynamics.R`.

#### Competitive communities

Competitive communities are those in which species only engage in intra- and interspecific competitive interactions. For these communities we generated random matrices with connectance $C = 0.2$, potentially asymmetric, and assumed that species display larger intra-specific than inter-specific competition, i.e. the interaction matrices are diagonally-dominant. We assumed that the system follows linear Lotka-Volterra dynamics given by the equation

$$\frac{dN_i}{dt} = \left( r_i + \sum_{j=1}^{S} \alpha_{ij} N_j \right) N_i, \quad \text{for } i = 1, 2, \ldots, S \qquad \text{(S1)}$$

where $S$ is the number of species in the system, $r_i$ is the intrinsic growth rate of species $i$, $\alpha_{ij}$ is the interaction strength coefficient from species $j$ to species $i$, and $N_i$ is the abundance of species $i$.

#### Food webs

We generated food webs following the niche model (Williams & Martinez, 2000), by randomly assigning a niche value in the interval $[0, 1]$ to each species in the community, related to its trophic position. Connectance of these food webs is the same as for the competitive and mutualistic communities, $C = 0.2$. Given these parameters (network connectance and niche values for all species), the food web structure is generated, very briefly, by assigning predator-prey links between predators with higher trophic position than their prey species. The dynamics of these food webs, similarly to the competitive communities, are simulated following the linear Lotka-Volterra model of eq. S1.

#### Mutualistic communities

Mutualistic interactions involve positive demographic effects for both of the interacting species, as in for example the plant-frugivore communities considered in this study. In population dynamics models such as the standard Lotka-Volterra formulation (eq. S1), the potential absence of density dependent self-limitation generates positive feedbacks of population growth and, therefore, biologically implausible dynamics (Hale & Valdovinos, 2021). Therefore, in mutualistic population dynamics, different strategies for constraining population growth have been proposed (Hale & Valdovinos, 2021). We implemented a population model for bipartite multi-species mutualistic communities with logistic population growth given by density-dependent benefits of mutualist interactions (García-Algarra *et al.*, 2014). This model, while relatively simple, can generate a rich array of dynamics, including asymptotically stable communities. Its formulation is

$$\frac{dN_i^a}{dt} = \left( r_i + \sum_{k=1}^{S_p} b_{ik} N_k^p - \left( \alpha_i + c_i \sum_{k=1}^{S_p} b_{ik} N_k^p \right) N_i^a \right) N_i^a, \quad \text{for } i = 1, 2, \ldots, S_a \tag{S2}$$

$$\frac{dN_j^p}{dt} = \left( r_j + \sum_{l=1}^{S_a} b_{jl} N_l^a - \left( \alpha_j + c_j \sum_{l=1}^{S_a} b_{jl} N_l^a \right) N_j^p \right) N_j^p, \quad \text{for } j = 1, 2, \ldots, S_p \tag{S3}$$

where the dynamics of the two guilds $a$ (plants) and $p$ (pollinators/seed dispersers) are given by equations S2 and S3. Terms $b_ij$ are the pairwise interaction strengths, $\alpha_i$ is a friction term that incorporates saturating benefits of mutualistic interactions, mediated by a proportionality constant $c_i$.

#### Obtaining dynamic net effects

We generated 100 interaction matrices for each combination of species richness and interaction type. For each of these replicates, we simulated numerically the dynamics of the system using the function `stode` from the package `rootSolve` v1.8.2.4 in R. We obtained the Jacobian matrix from the system at steady state (function `jacobian.full` from the `rootSolve` package), whose elements $A_{ij}$ can be interpreted as the direct effect of the average species $j$ individual on species $i$'s population growth rate. The net effects between each pair of species are the sum of direct and indirect effects between them, and are given by the negative of the inverse Jacobian matrix of the system (Novak *et al.*, 2016).

#### Communicability and dynamic net effects

In parallel to the net effects of the dynamical system, we obtained the pairwise communicability values $G_{ij}$ for each pair of species in each interaction matrix. For each network replicate, we obtained the Spearman correlation coefficient between the pairwise communicability values and the pairwise dynamic net effects, separating the species pairs between those that interact directly, and those that only do so through indirect paths. Our results show that pairwise communicability and dynamic net effects are strongly correlated 1) in all competitive communities, irrespective of their richness and type of pairwise link (direct or indirect), 2) in mutualistic communities, especially in less-rich communities and only for species interacting indirectly. In turn, in food webs and for directly interacting species in mutualistic communities, we found no or weak correlation between pairwise communicability and dynamic net effects (Fig S6).

These results reinforce the idea that the net demographic effect between pairs of species depends on the sign of the interactions in the community. In communities of a single interaction sign (competitive or mutualistic), the two metrics tend to be correlated; food webs however display positive (on predator species) and negative effects (on prey species), which potentially makes the patterns of aggregated net effects more complex, as these may include a mixture of positive and negative effects that cancel each other. The interpretation of the comparatively low correlation between communicability and dynamic net effects for species that interact directly in mutualistic communities is more nuanced, and likely related to the saturating effects of mutualistic interactions: net effects between species that interact directly may become neutral or negative at high densities, a process that is not captured by the communicability metric. Such neutral or negative effects between mutualistic partners, however, have only been documented empirically for patterns of high dominance of a single species, such as honey bees in agricultural systems (Rollin & Garibaldi, 2019).

When applying the communicability metric to empirical systems, therefore, one can be reasonably confident that the results obtained will be qualitatively similar to the dynamical net effects between species pairs for 1) communities with a single type of effect between each interacting species, 2) in mutualistic communities, when the number of species in the community is $< 60$. Further, in mutualistic communities, if there is a significant signal of density-dependent regulation, the estimation of communicability may differ considerably from that of dynamical net effects. In our study system, most of the local communities projected in our grid have richness values well below 60 (Fig. S7), so we assume that the insights from the communicability metric are reasonable first approximations to the propagation of effects in the system, given the absence of better empirical data. Further, the estimation of density-dependent demographic effects for diverse communities is challenging and out of the scope of our study, but as a first hypothesis, we posit that plant-frugivore interactions in our study have not generally reached a saturating benefit phase given the relatively low densities of bird species in most of the territory of Aotearoa New Zealand.

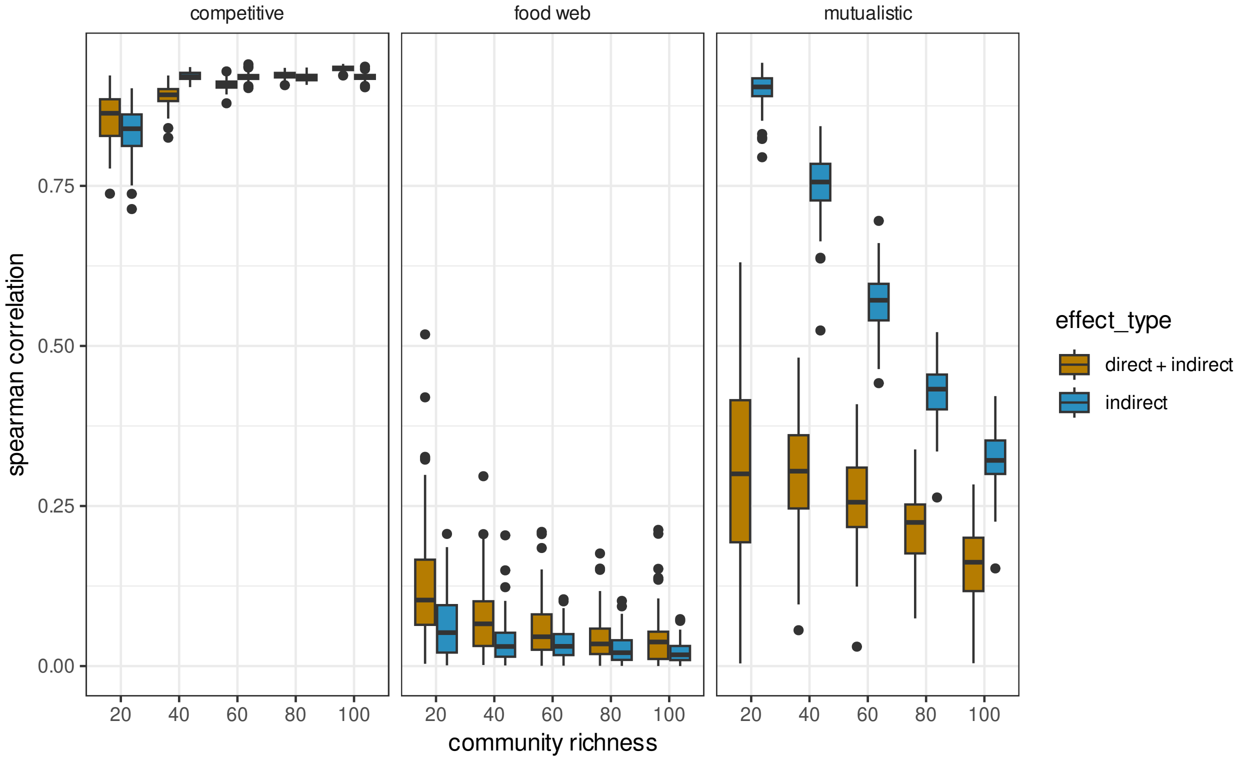


Fig. S6: Relationship between weighted communicability and dynamical net effects between pairs of species in simulated competitive communities, food webs, and mutualistic communities. Each panel represents a type of community, and each boxplot represents the distribution of correlation values from 100 network replicates. In the boxplots, the horizontal black lines represent the median, the lower and upper hinges correspond to the 25th and 75th percentiles, and the vertical lines extend to the largest/smallest value up to 1.5 times the interquartile range (distance between 25th and 75th percentiles).

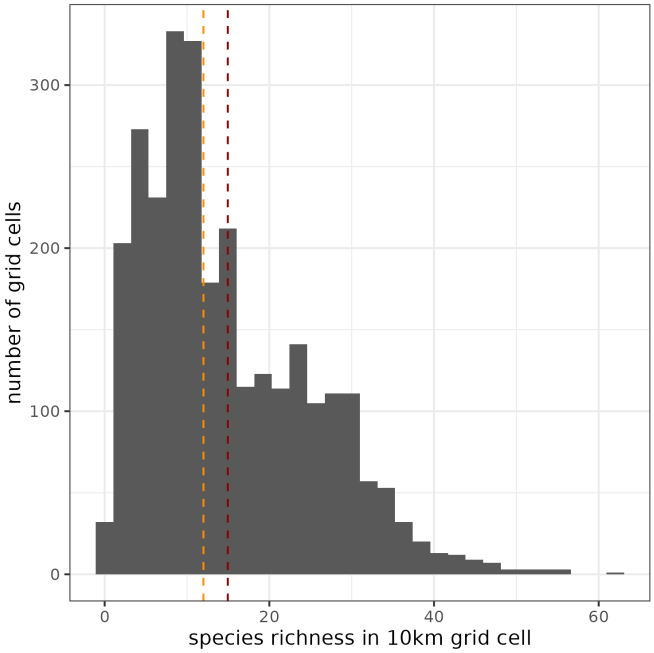


Fig. S7: Histogram of modelled species richness per 10km grid cell. Red dashed line represents the average of that distribution (15 species per cell). Orange dashed line represents the median (12 species per cell).

### Supplementary Section: Spatial network validation

We combined the modelled occurrences (Supplementary Section "Observed and predicted occurrences") with the metaweb of plant-frugivore interactions from the literature (Peralta *et al.*, 2020), to generate a local plant-frugivore network in each of the 3030 grid cells of our study area (Supplementary Section "Constructing the Aotearoa New Zealand plant-frugivore network"). Here we provide a validation of 1) the modelled species occurrences, and 2) the projected local networks.

#### Validation of modelled occurrences

We obtained the Area Under de ROC Curve (AUC, Jiménez-Valverde (2012)) of the modelled occurrences for each bird and plant species considered, using the `auc` function from the `pROC` package v1.18.5 in R. These were in all cases $> 0.8$, thus indicating very good agreement with the observed data (Fig. S8).

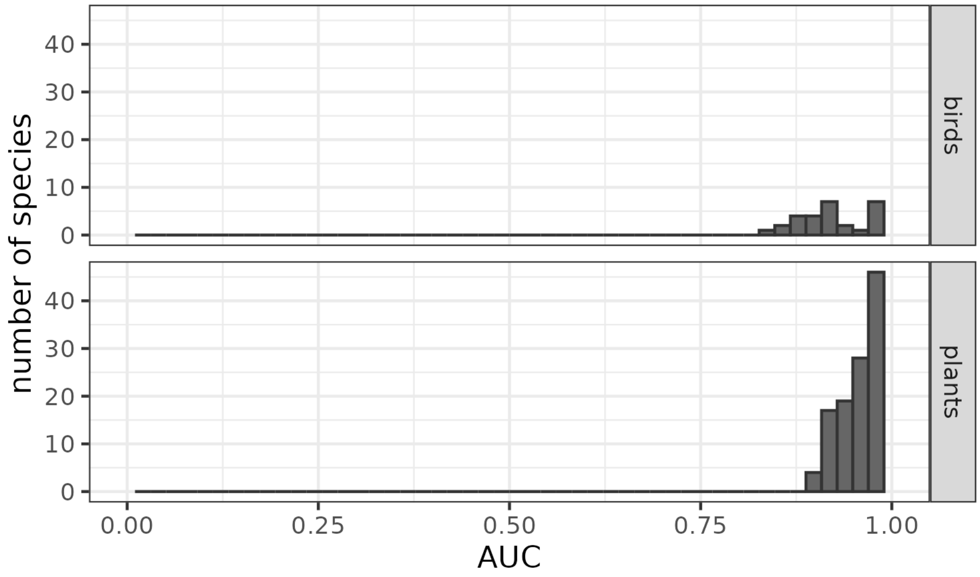


Fig. S8: Area under the ROC curve for the 102 plant and 22 bird species modelled in our networks. For each species, its AUC is calculated using the observed occurrences in the original pool of sites (940 TIER1 sites for birds, 1298 NVS sites for plants) and the modelled probabilities of occurrence from the JSDM as predictor.

#### Validation of local networks

For validating the local networks projected in the 10km cell grid, we gathered 16 plant-frugivore networks sampled across Aotearoa New Zealand (Fig. S9), from García *et al.* (2014) (9 networks), Burns (2013) (1 network), O'Donnell & Dilks (1994) (1 network), Williams & Karl (1996) (3 networks), Burrows (1994) (1 network), and MacFarlane *et al.* (2016) (3 networks). We denote $S(k)$ the species observed in the network at location $k$ (each of the 16 observed locations), $a_{i,j,k}$ the binary interaction between species $i, j \in S(k)$, which takes value $a_{i,j,k} = 1$ if an interaction between species $i$ and $j$ was observed empirically, and $a_{i,j,k} = 0$ otherwise. Likewise, we have the modelled probabilities of occurrence of species $i$ and $j$ given by the JSDM, and their probability of co-occurrence is $q_{i,j,k} = p_{i,k} * p_{j,k}$. Furthermore, the probability that species $i$ and $j$ are both present and interact at location $k$ is given by $z_{i,j,k} = q_{i,j,k} * m_{i,j}$, where $m_{i,j} = 1$ if the species interact in the compiled metaweb, and 0 otherwise. We first obtained the AUC values for the combinations $a_{i,j,k}, z_{i,j,k}$ at each location (Fig. S10). There was a high degree of variability, with 11 of 16 networks showing AUC values $\geq 0.6$, that is, from acceptable to good performance.

As a second step, to better understand the spatially varying nature of our projections, we asked if the observed network at location $k$ was better matched by the predicted network at $k$ than by the predicted network at any other location. In particular, we quantified the proportion of times than the AUC value for a location $k$ using the predicted network from $k$ was higher than using the predicted network from a different location. For example, if two of the 15 predicted networks from other locations were better predictors than the predicted network at $k$ for the observed network at $k$, this yields a relative frequency of $13/15 = 0.867$. The values for such relative frequencies of spatially explicit predictions, denoted $r$, were, again, highly variable (Fig. S11). Three networks showed particularly low values ($r < 0.3$), whereas nine out of 16 networks showed values of $r > 0.5$, which we deem as acceptable.

Overall, there are many sources of variability that affect the prediction of local interactions and its validation with independent data: from the species pool considered, to uncertainty associated with modelling species occurrences (Fig. S8), to uncertainty in predicting interaction presence/absence (Fig. S10) and its spatial variability (Fig. S11). These sources of uncertainty

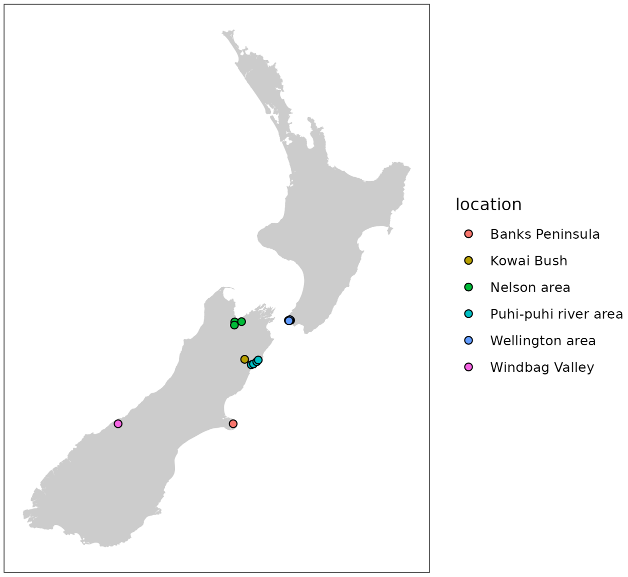


Fig. S9: Spatial location of the 16 plant-frugivore networks used to validate our projected networks. Six networks are located in the area of Wellington, four in the area of the Puhi-puhi river, and three in the area of Nelson.

impose practical limits to the ability to project ecological patterns across the territory. Our projections are, therefore, subject to such uncertainty, but we have shown that they provide generally acceptable predictions of species presences and absences, and of specific interactions when comparing with independently sampled datasets. A limitation to this validation exercise is the scarce number of independent local plant-frugivory networks that were available in our study area, and their spatial clustering.

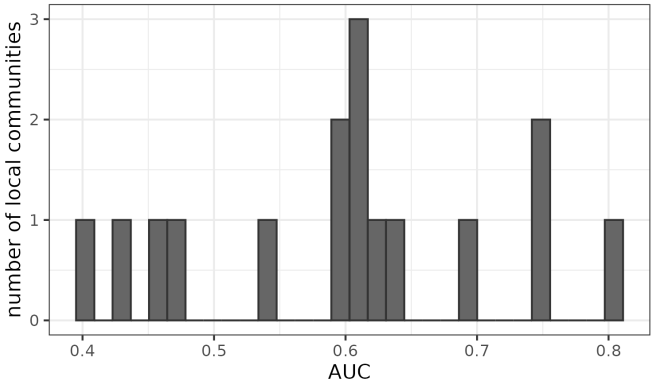


Fig. S10: AUC values for the combination of observed ($a_{i,j,k}$) and predicted ($p_{i,j,k}$) binary interactions for each of the 16 networks in Fig. S9.

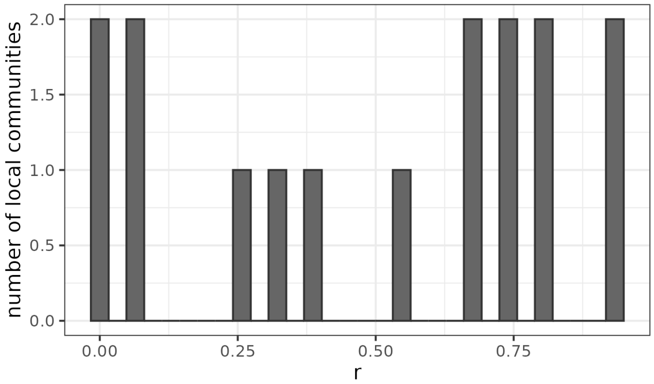


Fig. S11: For each of the 16 observed networks, relative frequencies of times when AUC values using its projected network at the same location give better matches than when using projected networks from other locations.

### Supplementary Section: Communicability of the Aotearoa New Zealand plant-frugivore network

To obtain the overall communicability of the national multilayer network required a full metacommunity structure in matrix form, i.e. a supra-adjacency matrix (Gravel *et al.*, 2016), which can then be decomposed per species or grid cell. The supra-adjacency matrix of the national multilayer network is of size $(3030 \times 124)^2$ (i.e. 3030 cells multiplied by 124 species on each dimension of the square matrix), which renders it unfeasible to analyze as a whole. Therefore, we calculated the communicability of the species in the studied area through a moving-window approach. We selected each of the 3030 cells sequentially and calculated the communicability $G_{i*}$ of local populations in that cell and a surrounding buffer of 30km, which represents a neighbourhood of 29 cells. This buffer distance is the maximum distance computationally feasible to calculate for our study area, and a preliminary test showed that the communicability rank of species did not vary significantly at increasing distances (i.e. the most important species are preserved; Figs. S12 and S13) because species separated by long indirect paths generally have weaker effects than those with shorter paths. This means that the spatial contributions to communicability are concentrated in a relatively close neighborhood of the focal area of the populations, which is consistent with the life history and movement patterns of the species considered here.

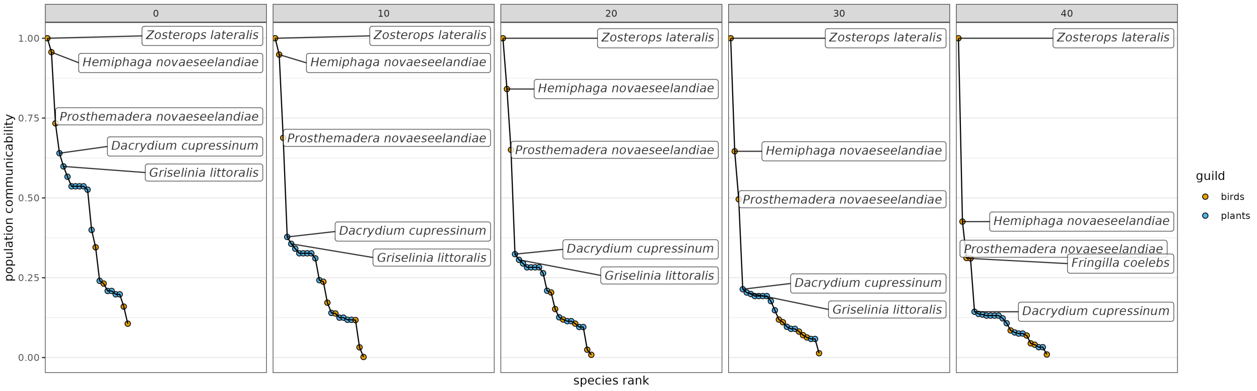


Fig. S12: Distributions of population communicability ranks from a focal cell with increasing numbers of neighboring cells included in the calculation, from cells in a buffer of 0km (i.e. only the focal cell) to 40km. Further distances, while theoretically computable, are computationally too costly to obtain, as the size of the supra-adjacency matrix grows exponentially.

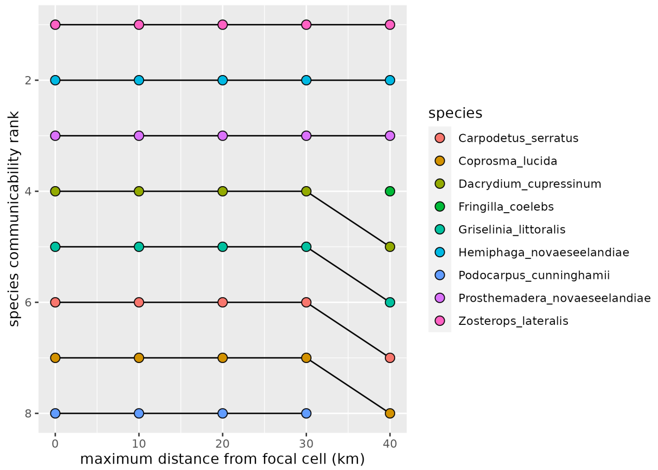


Fig. S13: Variations in communicability ranks of important species in a focal cell, as buffer distances for calculating communicability increase from 0km (i.e. only the focal cell to 40km. The three most important species (*Zosterops lateralis, Hemiphaga novaeseelandiae, Prosthemadera novaeseelandiae*) preserve their rank across increasing buffer distances.

### Supplementary Section: Correlation between species-level variables

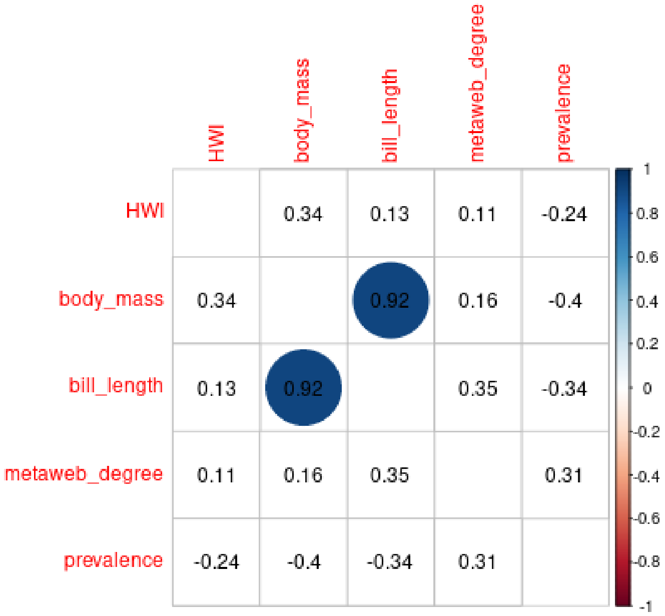


Fig. S14: Spearman correlations between species-level variables for birds. “HWI” is the species hand-wing index, the rest of the variables are self-explicative. Depicted are the correlation coefficients, and background circles are drawn for statistically significant correlations ($p < 0.05$).

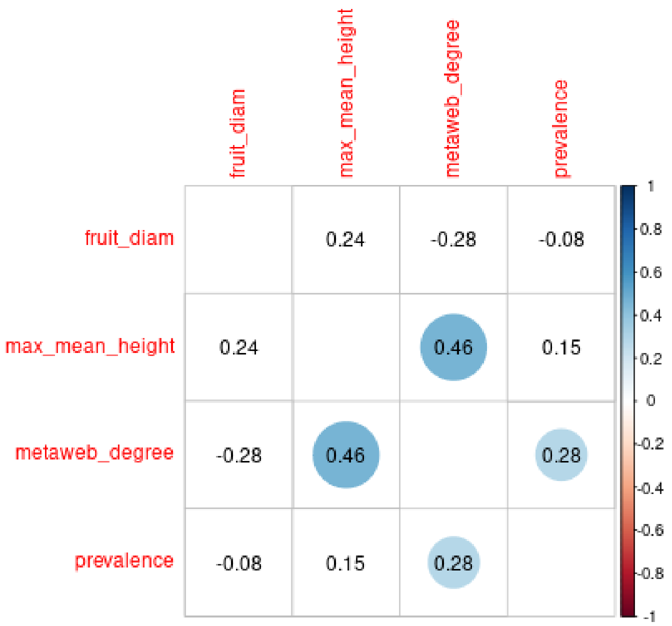


Fig. S15: Spearman correlations between species-level variables for plants. Depicted are the correlation coefficients, and background circles are drawn for statistically significant correlations ($p < 0.05$).

### Supplementary Section: Correlation between landscape variables

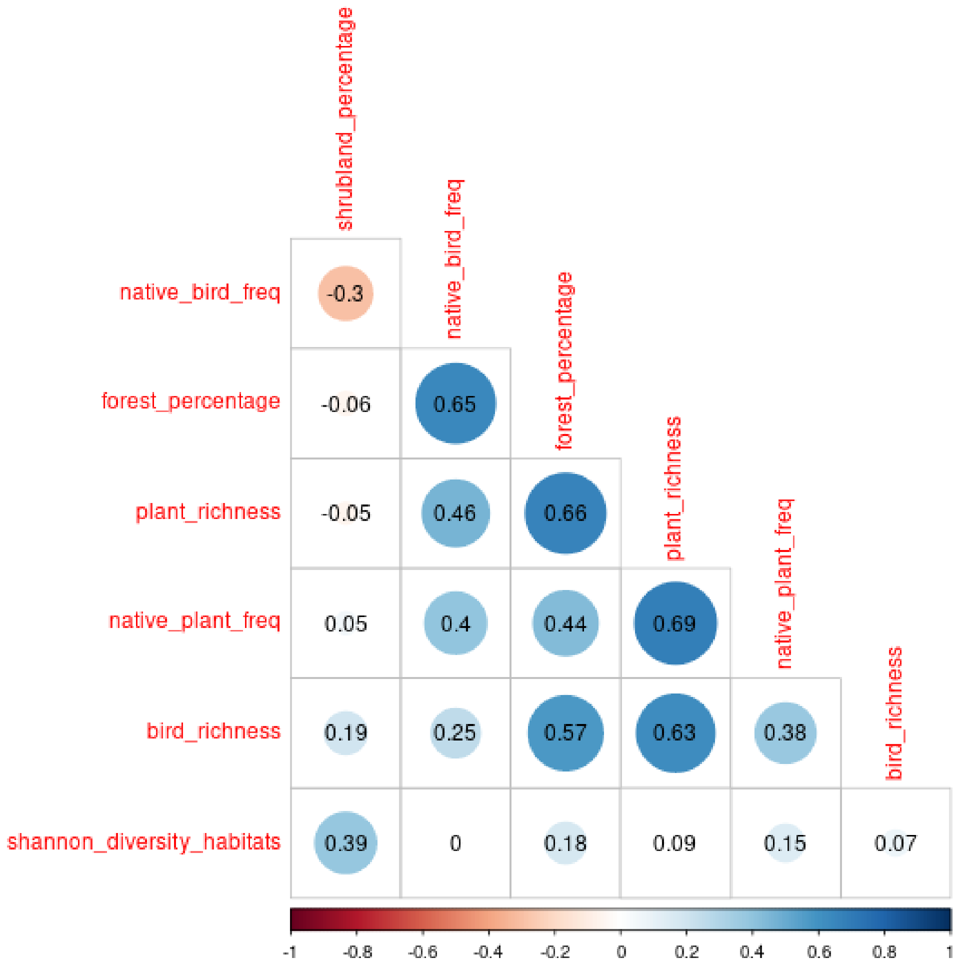


Fig. S16: Spearman correlations between cell-level variables. “native_bird_freq” and “native_plant_freq” are, respectively, the proportion of native bird/plant species from the total bird/plant richness in each cell. “forest_percentage” and “shrubland_percentage” are the percentage of these land uses in each cell’s area. “plant_richness” and “bird_richness” are the number of plant/bird species present in each cell. “shannon_diversity_habitats” is the Shannon diversity index applied to the proportions of land uses in each cell. Depicted are the correlation coefficients, and background circles are drawn for statistically significant correlations ($p < 0.05$).

### Supplementary Section: Supplementary regressions

Table S1: Coefficients of the generalised linear model (gamma distribution, log link) relating population-level communicability with local species degree, provenance, and species traits, for bird species. N = 22.

| | estimate | std.error | z | p.value |
|---|---|---|---|---|
| Intercept | -2.317 | 1.152 | -2.011 | 0.044 |
| status (Native) | 0.292 | 1.201 | 0.243 | 0.808 |
| degree | 0.754 | 0.292 | 2.584 | 0.010 |
| body mass (g) | -1.468 | 3.032 | -0.484 | 0.628 |
| status (Native):body mass (g) | 1.867 | 3.043 | 0.613 | 0.540 |

Table S2: Coefficients of the generalised linear model (gamma distribution, log link) relating population-level communicability with local degree, provenance, and species traits, for plant species. N = 100.

| | estimate | std.error | z | p.value |
|---|---|---|---|---|
| Intercept | -3.600 | 0.450 | -7.994 | $< 0.001$ |
| status (Native) | 1.435 | 0.460 | 3.121 | 0.002 |
| degree | 0.665 | 0.097 | 6.840 | $< 0.001$ |
| fruit diameter (mm) | -9.109 | 3.212 | -2.836 | 0.005 |
| max mean height (m) | 0.134 | 0.108 | 1.236 | 0.216 |
| status (Native):fruit diameter (mm) | 9.218 | 3.212 | 2.869 | 0.004 |

Table S3: Coefficientes of the generalised linear model relating landscape-level communicability with landscape characteristics from the neighbouring area (neighbouring cells in a 30km buffer zone). The expressive degrees of freedom of the spatial smoothing term is 28.14, with an F-value of 60.18 and a p-value $< 0.001$.

| | estimate | std.error | t | p.value |
|---|---|---|---|---|
| Intercept | 12.633 | 0.02 | 624.854 | $< 0.001$ |
| neighbor forest cover (%) | 1.643 | 0.04 | 41.315 | $< 0.001$ |
| neighbor shrubland cover (%) | 0.571 | 0.04 | 16.041 | $< 0.001$ |
| neighbor habitat diversity | -0.296 | 0.03 | -8.685 | $< 0.001$ |

### Supplementary Section: Variable importance estimation

In addition to estimating model coefficients and p-values from the statistical models discussed in the main text, we calculated the contribution of each factor to the marginal $R^2$ of the models, to complement their ecological interpretation. We performed this analysis with regression models that did not include the interaction term between body mass/fruit diameter and provenance, due to limitations of the methodology. Using these simpler models, here we report the percentage of the marginal $R^2$ explained by each predictor, obtained with the R package glmm.hp v0.1-0 (Lai *et al.*, 2022).

Table S4: Partitioning of marginal $R^2$ of the statistical model for bird species at the local level. Marginal $R^2$ of the model is 0.31.

| variable | percentage $R^2$ explained |
|---|---|
| provenance | 8.52 |
| body mass (g) | 16.85 |
| local network degree | 74.63 |

Table S5: Partitioning of marginal $R^2$ of the statistical model for plant species at the local level. Marginal $R^2$ of the model is 0.411.

| variable | percentage $R^2$ explained |
|---|---|
| provenance | 0.34 |
| fruit diameter (mm) | 1.14 |
| max mean height (m) | 13.07 |
| local network degree | 85.44 |

Table S6: Partitioning of marginal $R^2$ of the statistical model for bird species at the country level. Marginal $R^2$ of the model is 0.54.

| variable | percentage $R^2$ explained |
|---|---|
| provenance | 4.16 |
| body mass (g) | 6.18 |
| prevalence | 56.42 |
| metaweb degree | 33.24 |

Table S7: Partitioning of marginal $R^2$ of the statistical model for plant species at the country level. Marginal $R^2$ of the model is 0.7.

| variable | percentage $R^2$ explained |
|---|---|
| provenance | 18.19 |
| fruit diameter (mm) | 1.24 |
| max mean height (m) | 3.34 |
| prevalence | 50.34 |
| metaweb degree | 26.9 |

### Supplementary Figures

The following figures were generated with the R package ggeffects v1.8.5.

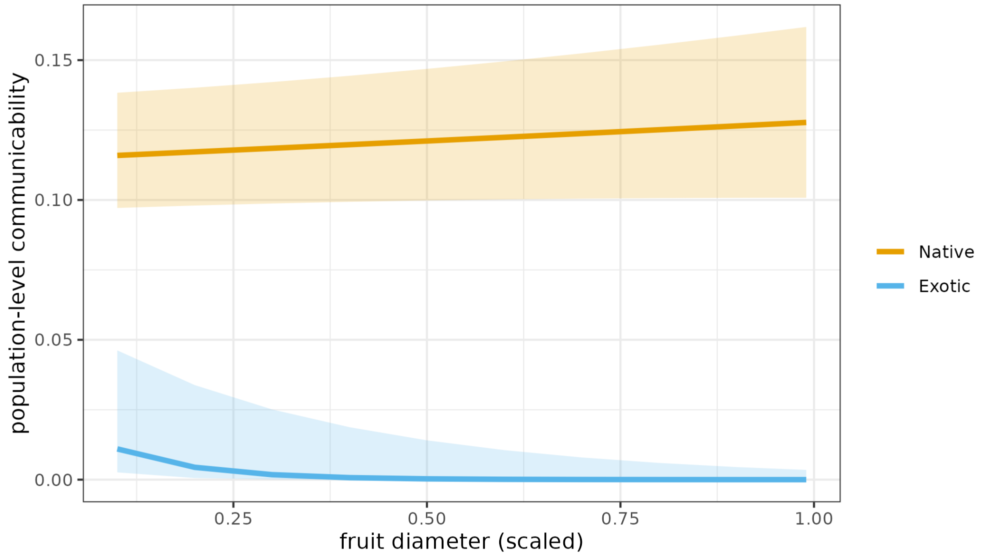


Fig. S17: Effect of fruit diameter on population-level communicability for plant species, accounting for the interaction with species provenance. Shaded areas represent 95% confidence intervals.

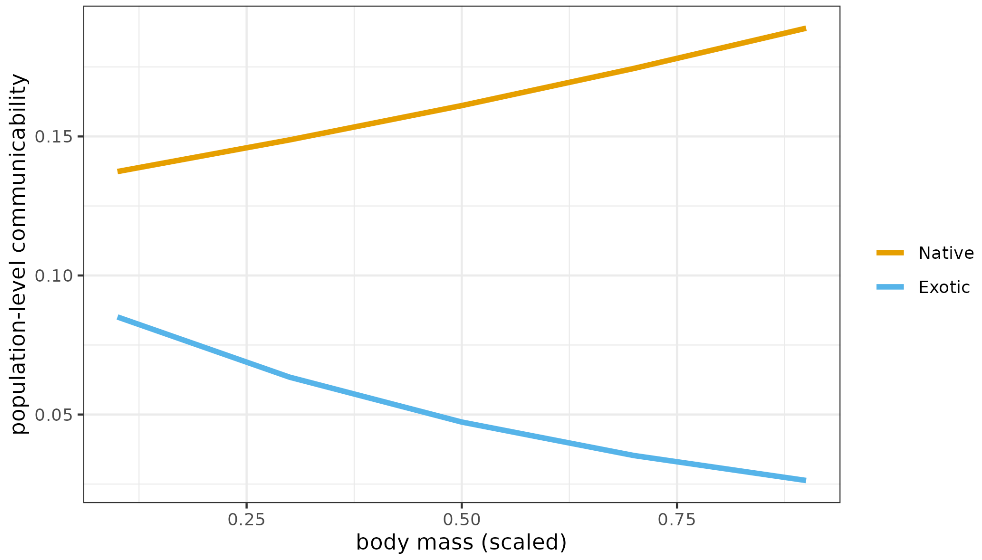


Fig. S18: Effect of body mass on population-level communicability for bird species, accounting for the interaction with species provenance. Confidence intervals not shown for visibility, as they fully overlap.

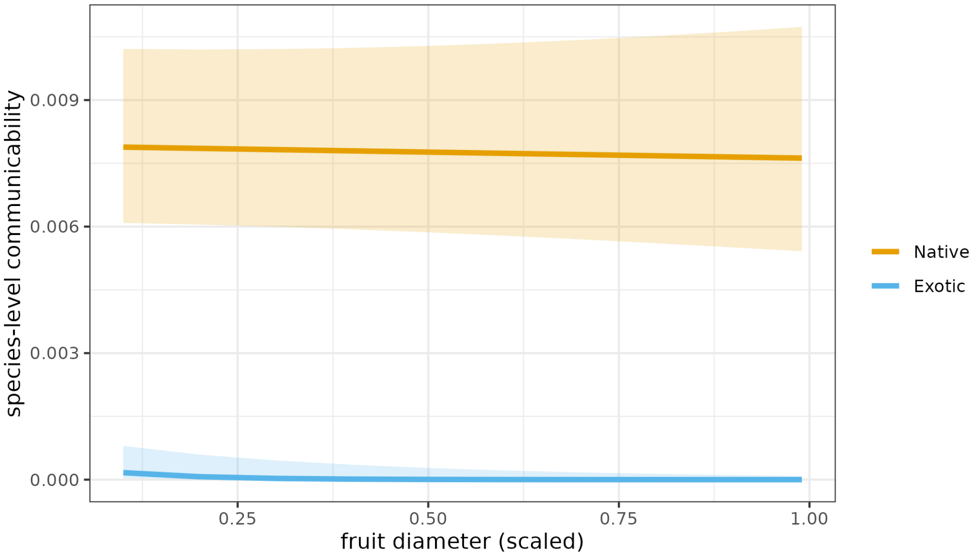


Fig. S19: Effect of fruit diameter on species-level communicability for plant species, accounting for the interaction with species provenance. Shaded areas represent 95% confidence intervals.

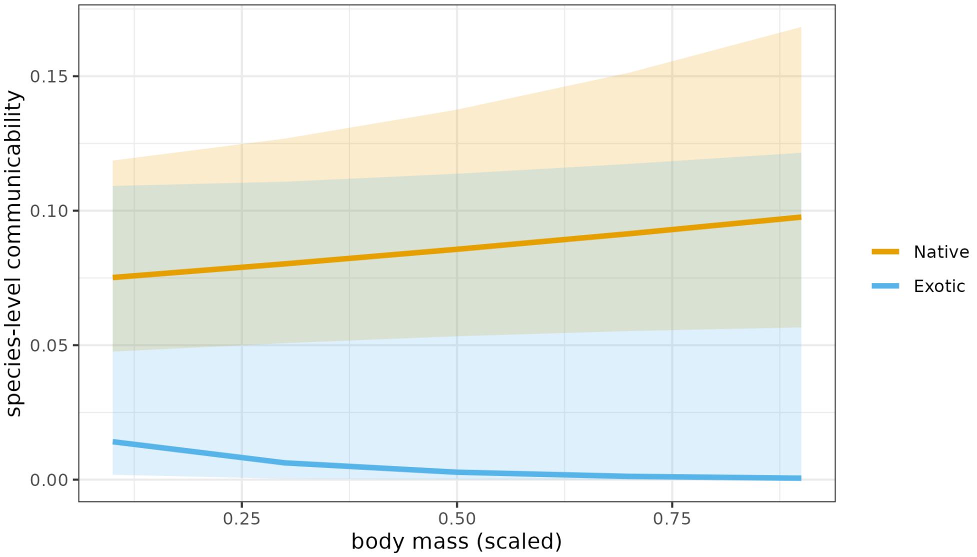


Fig. S20: Effect of body mass on species-level communicability for bird species, accounting for the interaction with species provenance. Shaded areas represent 95% confidence intervals.

### Supplementary References

Burns, K. (2013). What causes size coupling in fruit–frugivore interaction webs? *Ecology*, 94, 295–300.

Burrows, C. J. (1994). Fruits, seeds, birds and the forests of banks peninsula. *New Zealand Natural Sciences*, 21, 87–108.

García, D., Martínez, D., Stouffer, D. B. & Tylianakis, J. M. (2014). Exotic birds increase generalization and compensate for native bird decline in plant–frugivore assemblages. *Journal of Animal Ecology*, 83, 1441–1450.

García-Algarra, J., Galeano, J., Pastor, J. M., Iriondo, J. M. & Ramasco, J. J. (2014). Rethinking the logistic approach for population dynamics of mutualistic interactions. *Journal of theoretical biology*, 363C, 332–343.

García-Callejas, D., Molowny-Horas, R., Araújo, M. B. & Gravel, D. (2019). Spatial trophic cascades in communities connected by dispersal and foraging. *Ecology*, 100, e02820.

Gravel, D., Massol, F. & Leibold, M. A. (2016). Stability and complexity in model meta-ecosystems. *Nature communications*, 7, 12457.

Hale, K. R. S. & Valdovinos, F. S. (2021). Ecological theory of mutualism: Robust patterns of stability and thresholds in two-species population models. *Ecology and Evolution*, 11, 17651–17671.

Jiménez-Valverde, A. (2012). Insights into the area under the receiver operating characteristic curve (AUC) as a discrimination measure in species distribution modelling. *Global Ecology and Biogeography*, 21, 498–507.

Lai, J., Zou, Y., Zhang, S., Zhang, X. & Mao, L. (2022). glmm.hp: an r package for computing individual effect of predictors in generalized linear mixed models. *Journal of Plant Ecology*, 15, 1302–1307.

MacFarlane, A., Kelly, D. & Briskie, J. (2016). Introduced blackbirds and song thrushes: Useful substitutes for lost mid-sized native frugivores, or weed vectors? *New Zealand Journal of Ecology*, 40, 80–87.

Novak, M., Yeakel, J. D., Noble, A. E., Doak, D. F., Emmerson, M., Estes, J. A., Jacob, U.,

Tinker, M. T. & Wootton, J. T. (2016). Characterizing species interactions to understand press perturbations: What is the community matrix? *Annual Review of Ecology, Evolution, and Systematics*, 47, 409–432.

O'Donnell, C. F. & Dilks, P. J. (1994). Foods and foraging of forest birds in temperate rainforest, south westland, new zealand. *New Zealand Journal of Ecology*, 18, 87–107.

Peralta, G., Perry, G. L. W., Vázquez, D. P., Dehling, D. M. & Tylianakis, J. M. (2020). Strength of niche processes for species interactions is lower for generalists and exotic species. *Journal of Animal Ecology*, 89, 2145–2155.

Poisot, T. & Gravel, D. (2014). When is an ecological network complex? Connectance drives degree distribution and emerging network properties. *PeerJ*, 2, e251.

Rollin, O. & Garibaldi, L. A. (2019). Impacts of honeybee density on crop yield: A meta-analysis. *Journal of Applied Ecology*, 56, 1152–1163.

White, J. W., Rassweiler, A., Samhouri, J. F., Stier, A. C. & White, C. (2014). Ecologists should not use statistical significance tests to interpret simulation model results. *Oikos*, 123, 385–388.

Williams, P. A. & Karl, B. J. (1996). Fleshy fruits of indigenous and adventive plants in the diet of birds in forest remnants, nelson, new zealand. *New Zealand journal of ecology*, 20, 127–145.

Williams, R. J. & Martinez, N. D. (2000). Simple rules yield complex food webs. *Nature*, 404, 180–3.
